# Ultrasensitive Molecular Imaging of Mucosal Inflammation Using Leucocyte-Mimicking Particles Targeted to MAdCAM-1

**DOI:** 10.1101/771659

**Authors:** Antoine P. Fournier, Sara Martinez de Lizarrondo, Adrien Rateau, Axel Gerard-Brisou, Maximilian J. Waldner, Markus F. Neurath, Denis Vivien, Fabian Docagne, Maxime Gauberti

## Abstract

Mucosal tissues line the digestive, respiratory, urinary, mammary and reproductive tracts and play critical roles in health and disease as the primary barrier between the external world and the inner body. Clinical evaluation of mucosal tissues is currently performed using endoscopy, such as ileocolonoscopy for the intestinal mucosa, that causes significant patient discomfort and can lead to organ damage. Here, we developed a new contrast agent for molecular magnetic resonance imaging (MRI) that is targeted to mucosal vascular addressin cell adhesion molecule 1 (MAdCAM-1), an adhesion molecule overexpressed by inflamed mucosal tissues. We investigated the diagnostic performance of molecular MRI of MAdCAM-1 to detect mucosal inflammation in several models of acute and chronic intestinal inflammation in mice. We demonstrated that molecular MRI of MAdCAM-1 reveals disease activity and can evaluate the response to inflammatory treatments along the whole intestinal mucosa in clinically relevant models of inflammatory bowel diseases. We also provide evidence that this new technique can detect low, subclinical levels of mucosal inflammation. Molecular MRI of MAdCAM-1 has thus potential applications in early diagnosis, longitudinal follow-up and therapeutic response monitoring in diseases affecting mucosal tissues, such as inflammatory bowel diseases.

**One Sentence Summary:** Molecular magnetic resonance imaging allows non-invasive evaluation of mucosal inflammation in clinically relevant experimental models.

## INTRODUCTION

Mucosal tissues line the digestive, respiratory, urinary, mammary and reproductive tracts at the interface between the microbiota and the inner body. They are involved in a very wide range of diseases from infection with pathogens, to autoimmune diseases and cancer. Although their total surface area exceeds 30 square meters in humans(*1*), the small thickness of mucosal tissues prevents their reliable evaluation by non-invasive imaging such as magnetic resonance imaging (MRI), computed tomography and positron emission tomography. Therefore, evaluation of inner mucosal tissues in patients is currently based on visual inspection by endoscopy, such as bronchoscopy for the lung, cystoscopy for the urinary bladder and ileocolonoscopy for the lower end of the gastrointestinal tract. Importantly, histological sampling performed during these endoscopic procedures can only explore a very limited part of the involved mucosal tissues, thereby putting the patients at risk of underdiagnosis of potentially life-threatening diseases. These limitations significantly impair clinician ability to detect early mucosal lesions, to spatially map mucosal damages in acute diseases and to accurately follow chronically ill patients.

Among the diseases affecting mucosal tissues, inflammatory bowel diseases (IBD) have become a major medical burden since they are the second most common chronic inflammatory disease worldwide and their prevalence has been increasing over the past decades in both adult and pediatric populations(*2*). Crohn’s disease and ulcerative colitis are the two principal subtypes of IBD and involve chronic immunological response in the gastrointestinal mucosa. Ileocolonoscopy with biopsies is still considered the gold standard for initial diagnosis, disease activity monitoring and assessment of therapeutic response(*3*). Unfortunately, this procedure is costly, causes significant patient discomfort, can lead to intestinal perforation and is limited to the evaluation of the lower part of the gastrointestinal mucosa (rectum, colon and terminal ileum)(*4*). Alternatives, such as MRI, can explore the entire gastrointestinal tract, detect severe intestinal lesions and complications but cannot detect isolated mucosal inflammation nor ascertain mucosal healing after treatment(*5*). In this context, there is a need for a new diagnostic tool allowing non-invasive and sensitive assessment of the inflammatory status of the mucosa along the whole gastrointestinal tract.

At the molecular level, mucosal vascular addressin cell adhesion molecule 1 (MAdCAM-1) represents an interesting biomarker for mucosal inflammation(*6*). Indeed, inflammation in mucosal tissue is associated with several phenotypical changes including activation of the endothelial cells of the lamina propria(*7*). Upon activation, these endothelial cells overexpress MAdCAM-1, which is responsible for the trafficking of leucocytes in these tissues(*6*). Accordingly, MAdCAM-1 is significantly overexpressed in active IBD and vedolizumab, a monoclonal antibody directed against the ligand of MAdCAM-1 (the integrin α4β7), is currently used as a treatment for both Crohn disease and ulcerative colitis. Non-invasive detection of MAdCAM-1 could therefore be used to assess the inflammatory status of the intestinal mucosa. Importantly, inflammation is a hallmark of mucosal damage and is found not only in autoimmune but also in other disorders affecting mucosal tissues such as cancers(*8*).

Recently, we and others reported the use of a new family of contrast agents for molecular MRI based on micro-sized particles of iron oxide (MPIO) that displays a dramatically higher sensitivity than classically used nano-sized particles(*9–12*). Thanks to an artefact known as the blooming effect, MPIO appear around 50 times larger than their actual size on iron sensitive T2*-weighted images (*12*). This characteristic of MPIO makes them particularly interesting for imaging mucosal tissues, since these tissues measure only a few hundred micrometers of thickness in physiological conditions and are thus hardly visible on unenhanced MRI. In the present study in mice, we developed a new MPIO-based contrast agent for MRI targeted to MAdCAM-1 (thereafter named MPIO-αMAdCAM-1). In three different models of intestinal inflammation, including lipopolysaccharides (LPS) induced endothelial activation, dextran sodium sulfate (DSS) induced colitis and trinitrobenzene sulfonic acid (TNBS) induced colitis(*13*), we demonstrated that this contrast agent allows detection of mucosal inflammation, quantification of disease activity and assessment of the mucosal response to anti-inflammatory treatments.

## RESULTS

### Effective bioconjugation of anti-MAdCAM-1 monoclonal antibodies to tosylactived MPIO

We produced MPIO-αMAdCAM-1 by covalently coupling micro-sized particles of iron oxide to monoclonal antibodies targeted against MAdCAM-1 (**Figure 1A**). We chose covalent coupling and monoclonal antibodies to obtain the highest bonding force between the MPIO and its mucosal target, with the aim of stabilizing the particle after binding and thereby allowing imaging during a sufficiently large time window. We confirmed effective bioconjugation of the monoclonal antibodies on the surface of MPIO by both flow cytometry (**Figure S1**) and confocal microscopy (**Figure S2**). Both methods demonstrated that, after conjugation, anti-MAdCAM-1 monoclonal antibodies can be detected on the surface of MPIO-αMAdCAM-1 using fluorescent secondary antibodies. Importantly, a sonication step was added during the preparation of the conjugated MPIO in order to obtain an almost monodisperse suspension of particles as evidenced on confocal microscopy images (**Figure S2**). This sonication step avoided the presence of aggregate that can induce false positive findings by plugging into the microcirculation after intravenous injection(*14*).

**Figure 1.**
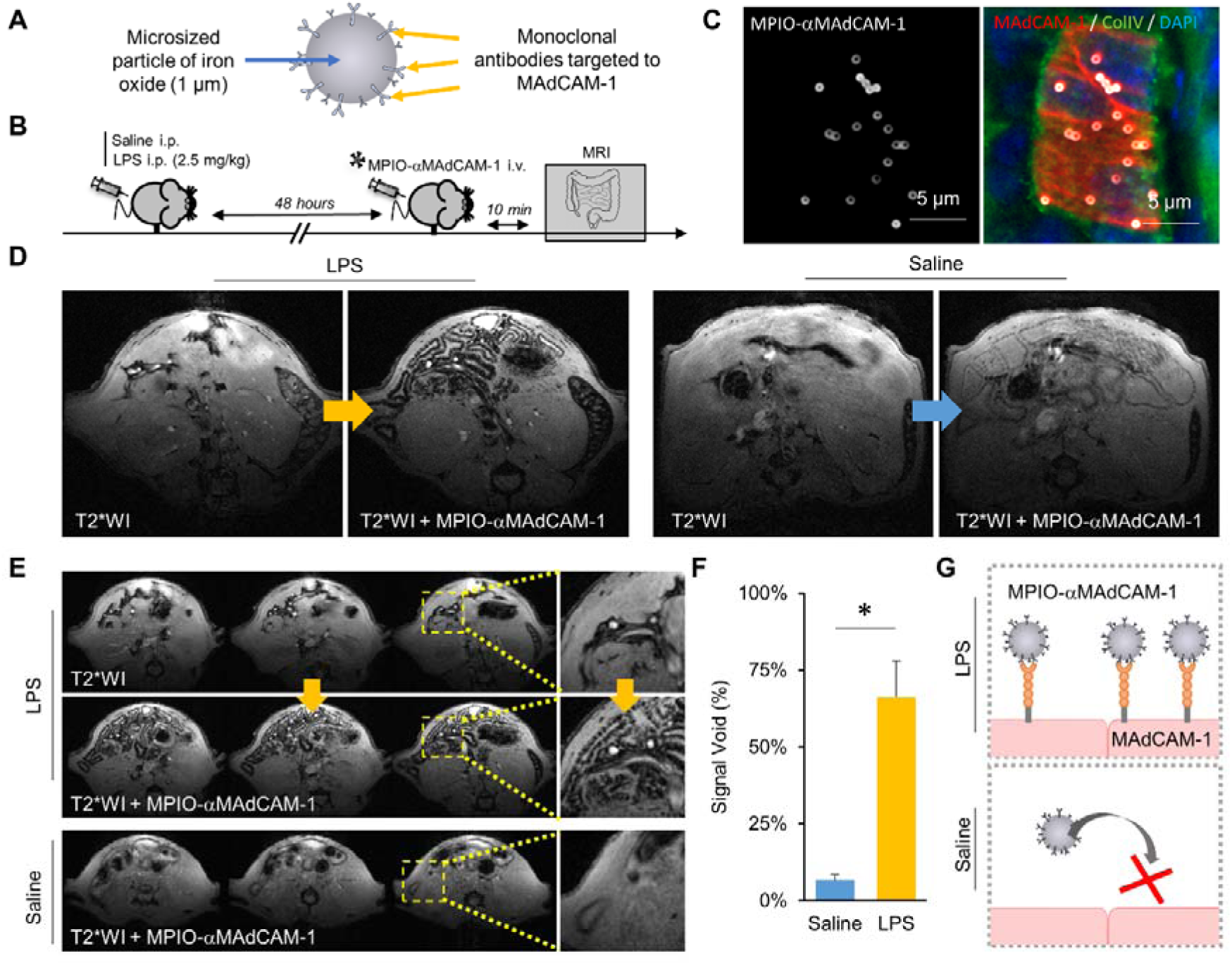
MPIO-αMAdCAM-1 binds to the activated endothelium of the gastrointestinal tract. (A) Schematic representation of the structure of MPIO-αMAdCAM-1. (B) Schematic representation of the experimental procedure. (C) Representative immunofluorescence images of MPIO-αMAdCAM-1 on the surface of a MAdCAM-1 positive vessel in a mouse that received LPS 48 hours before MPIO-αMAdCAM-1 injection. (D) Representative T2*-weighted images of the abdomen of mice treated with LPS (left) or saline (right) both before and after intravenous injection of MPIO-αMAdCAM-1. (E) Representative multi-slice T2*-weighted images of the abdomen of a mouse treated with LPS (upper part) both before and after intravenous injection of MPIO-αMAdCAM-1 and of a mouse treated with saline (lower part) including magnification on some intestinal loops presented on the left. Note that no contrast change occurs in the kidney after injection of the MPIO-αMAdCAM-1 in line with the lack of renal expression of MAdCAM-1. (F) Corresponding quantification (n=5 per group). (G) Schematic representation of the findings. *p<0.05 vs the indicated group.

### MPIO-**α**MAdCAM-1 binds specifically to the activated endothelium of the gastrointestinal tract

Our next objective was to demonstrate that MPIO-αMAdCAM-1 can bind activated endothelial cells in the intestinal mucosa *in vivo* after intravenous injection. To induce endothelial activation in the intestinal mucosa of mice, we performed an intraperitoneal injection of LPS (**Figure 1B**). Immunohistological analyses performed 48 hours thereafter confirmed overexpression of MAdCAM-1 in the lamina propria of the mucosa of the ileum of LPS treated compared to control saline treated mice (**Figure S3**). To determine whether systemically administered MPIO-αMAdCAM-1 can bind intestinal MAdCAM-1, we performed immunohistological analyses of the ileum 10 minutes after intravenous injection of MPIO-αMAdCAM-1 in LPS treated mice. In these mice, numerous MPIO-αMAdCAM-1 were bound to MAdCAM-1 expressing vessels in the lamina propria of the ileum (**Figure 1C**). Importantly, ∼99% of MPIO-αMAdCAM-1 were located on the inner surface of MAdCAM-1 expressing vessels with no apparent extravasated particle (**Figure S4**). The next step was to demonstrate that MPIO-αMAdCAM-1 bound to the endothelial cells of the intestinal mucosa can be detected by MRI. To this aim, we performed T2*-weighted MRI of the intestines both before and 10 minutes after intravenous injection of MPIO-αMAdCAM-1 (48 hours after intraperitoneal injection of LPS or saline). Post-contrast images in LPS-treated mice revealed MPIO-αMAdCAM-1 induced signal voids in the mucosa of the imaged intestinal loops (**Figure 1D**, **Figure S5 and Movie S1**). These signal void created a black lining at the center of the intestinal loops on post-contrast T2*-weighted images, in line with the expected distribution of MAdCAM-1 in intestinal mucosa. Unlike LPS-treated mice, saline treated mice presented only minimal MRI signal change on post-contrast images (**Figure 1D-G**).

We confirmed that the observed MRI signal changes were due to the magnetic susceptibility effect of the MPIO-αMAdCAM-1 bound to MAdCAM-1 expressed by the activated endothelium of the inflamed mucosa by showing that i) LPS-treated mice that received control, untargeted MPIO-αIgG (i.e. unable to bind MAdCAM-1) presented no detectable signal change on post-contrast images (**Figure 2A-D**) and ii) LPS-treated mice that received a blocking dose of anti-MAdCAM-1 antibody to saturate the binding sites before injection of the MPIO-αMAdCAM-1 presented significantly less signal changes on post-contrast images compared to control IgG-treated mice (**Figure 2E-H**). Longitudinal imaging after a single injection of MPIO-αMAdCAM-1 revealed that MPIO-αMAdCAM-1 progressively unbind from the inflamed endothelium with no residual contrast evidenced after 4 hours (**Figure S6**), again confirming that the MPIO-αMAdCAM-1 responsible for the observed signal voids are intraluminal (i.e. not extravasated) and capable of recirculate after unbinding. Altogether these results demonstrate that MPIO-αMAdCAM-1 binds specifically to the activated endothelium of the intestinal mucosa with minimal binding in control mice and without observable particle extravasation.

**Figure 2.**
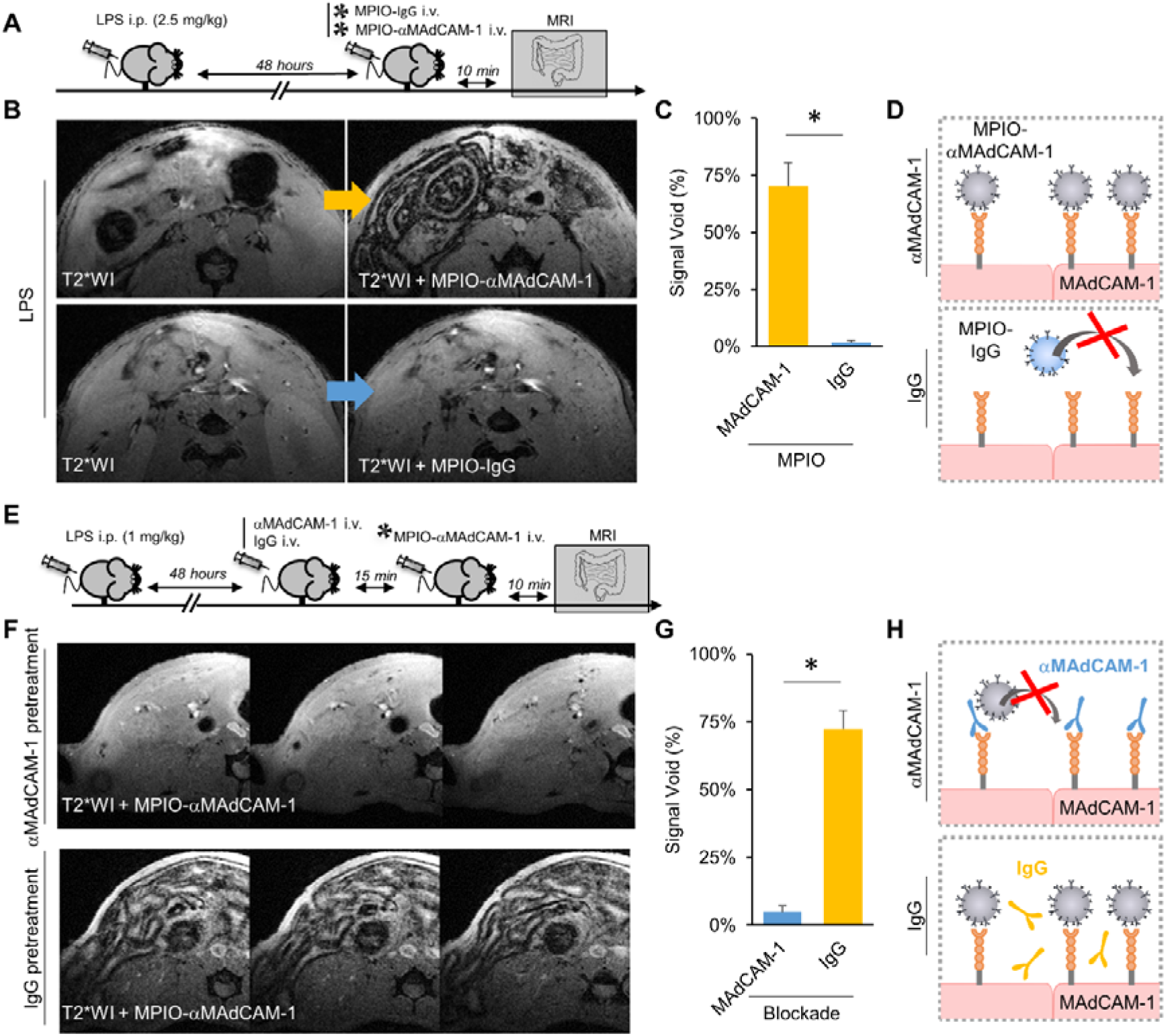
MPIO-αMAdCAM-1 binding is specific. (A) Schematic representation of the first experimental procedure. (B) Representative T2*-weighted images of the abdomen of mice treated with LPS both before and after intravenous injection of either MPIO-αMAdCAM-1 (upper part) or MPIO-IgG (lower part). (C) Corresponding quantification (n=5 per group). (D) Schematic representation of the findings from panels (A) to C). (E) Schematic representation of the second experimental procedure. (F) Representative multi-slice T2*-weighted images of the abdomen of mice treated with LPS and that received either antibodies against MAdCAM-1 (αMAdCAM-1 pretreatment, upper part) or control antibodies (IgG pretreatment, lower part) 15 minutes before MPIO-αMAdCAM-1 injection. (G) Corresponding quantification (n=5 per group). (H) Schematic representation of the findings from panels (E) to (G). *p<0.05 vs the indicated group.

### MPIO-**α**MAdCAM-1 enhanced MRI reveals mucosal inflammation after DSS-induced acute colitis

Then, we wanted to investigate the value of MPIO-αMAdCAM-1 enhanced imaging to reveal mucosal inflammation in more clinically relevant experimental models. The first model of colitis that we used was the DSS-model of acute colitis(*15*). DSS induces intestinal inflammation by disrupting the intestinal epithelial monolayer lining, leading to the entry of luminal bacteria and associated antigens into the mucosa, triggering an immune response. Mice were fed during five days with different concentrations of DSS in the drinking water (from 0% to 4%, **Figure 3A**) to induce different levels of colitis severity. After five days of treatment, we performed axial imaging of the descending colon including T2-weighted sequences to reveal mucosal thickening and submucosal edema, two morphological imaging biomarkers of intestinal inflammation, and T2*-weighted sequences both before and after MPIO-αMAdCAM-1 injection. Clinical symptoms of colitis as well as submucosal edema in the descending colon on T2-weighted images were evident in mice treated with 2% and 4% of DSS (**Figure 3B-C** and **Figure S7**), confirming active disease. Post-contrast images after intravenous injection of MPIO-αMAdCAM-1 revealed significant overexpression of MAdCAM-1 in the mucosa of those mice (**Figure 3D-E**). In contrast, control, untargeted MPIO-IgG failed to induce significant contrast change, confirming the specificity of the contrast agent in this model (**Figure S8**). Interestingly, mice that received 1% of DSS did not present clinical symptoms of colitis and their unenhanced T2-weighted images were undistinguishable from control mice. However, MPIO-αMAdCAM-1 enhanced MRI revealed a higher level of mucosal inflammation in these mice compared to control mice, suggesting that molecular MRI of MAdCAM-1 can reveal subclinical levels of intestinal inflammation (**Figure 3D-E**).

**Figure 3.**
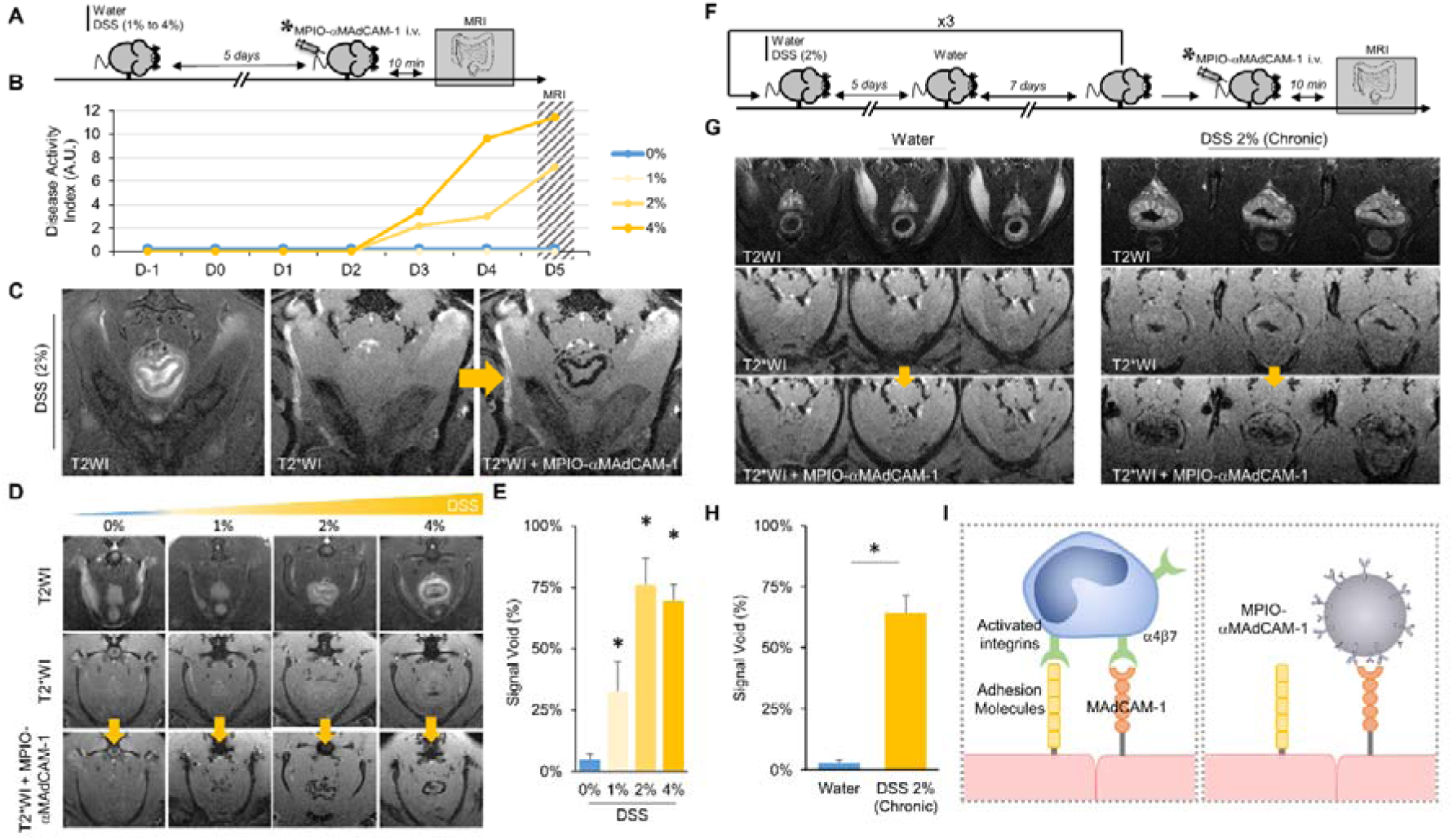
MPIO-αMAdCAM-1 enhanced MRI reveals mucosal inflammation after DSS-induced acute and chronic colitis. (a) Schematic representation of acute colitis induction using DSS at different concentrations (from 0% to 4%). (b) Mean disease activity index during the 5 days of DSS treatment (n=5 per group). (c) Representative T2-weighted, T2*-weighted and T2*-weighted after injection of MPIO-αMAdCAM-1 images in a mouse that received 2% DSS for 5 days. (d) Representative T2-weighted, T2*-weighted and T2*-weighted after injection of MPIO-αMAdCAM-1 images in mice that received different concentrations of DSS for 5 days. (e) Corresponding quantification (n=5 per group). *p<0.05 vs the 0% DSS group (water). (f) Schematic representation of chronic colitis induction using 2% DSS in three 5-day cycles. (g) Representative T2-weighted, T2*-weighted and T2*-weighted after injection of MPIO-αMAdCAM-1 images in control (water) and chronic colitis mice (DSS 2%). (h) Corresponding quantification (n=5). *p<0.05 vs the 0% DSS group (water). (i) Schematic representation of the behavior of MPIO-αMAdCAM-1 in the colitis model, that act as leucocyte mimicking particles.

We also used the acute DSS model to better characterize the kinetic of MPIO-αMAdCAM-1 binding after intravenous injection (**Figure S9 and Movie S2)**. We performed longitudinal fast T2*-weighted imaging (4s of temporal resolution) to detect MPIO-αMAdCAM-1 in the vasculature (caudal vein) in the colonic mucosa (to measure specific binding) and in paraspinal muscles (to measure nonspecific binding). Our results indicate that the plasmatic half-life of MPIO-αMAdCAM-1 is very short (< 1 min) with no detectable vascular signal after 120 seconds. Accordingly, MPIO-αMAdCAM-1 binding in the colonic mucosa reached its maximum ∼80 seconds after intravenous injection and remained stable thereafter during the period of observation (140 seconds). No detectable signal change was observed in the paraspinal muscles.

### MPIO-**α**MAdCAM-1 enhanced MRI reveals mucosal inflammation after DSS-induced chronic colitis

Imaging of chronic colitis represents a daily challenge in clinical practice since currently available imaging methods cannot easily discriminate chronic morphological changes such as fibrosis from active mucosal inflammation(*16*). In this context, the ability to directly reveal endothelial activation in chronic inflammatory lesions would be of great interest to ascertain the existence of an active inflammatory process. To investigate whether MPIO-αMAdCAM-1 enhanced imaging can reveal mucosal inflammation in chronic colitis, we induced chronic colitis in mice using DSS as described elsewhere(*13*). Control mice received water instead of 2% DSS. After three cycles of 5 days 2% DSS or water administration, we performed T2-weighted imaging and T2*-weighted imaging both before and after MPIO-αMAdCAM-1 injection (**Figure 3F**). Morphological T2-weighted images revealed thickening and increased signal of the colonic mucosa in this DSS-induced chronic colitis model compared to control mice (**Figure 3G**). Notably, in this model of chronic colitis, no submucosal edema was evident on morphological images. In contrast, MPIO-αMAdCAM-1 enhanced imaging revealed significant overexpression of MAdCAM-1 in the mucosa (**Figure 3G-H**), confirming active inflammation. These data demonstrate that molecular MRI of MAdCAM-1 unmasks active inflammation in chronic lesions of the intestinal tract. Overall and in line with our current knowledge of immune cell trafficking in IBD(*6*), MPIO-αMAdCAM-1 act as leucocyte mimicking particles and display increased binding to the endothelium of the mucosa in both acute and chronic colitis (**Figure 3I**).

### Assessing therapeutic response in acute and chronic colitis using MPIO-**α**MAdCAM-1 enhanced imaging

The current aim of IBD therapies is to achieve complete arrest of the inflammatory process(*17, 18*), an endpoint associated with better long-term outcome than clinical remission alone(*19*). Assessment of therapeutic response, mucosal and histological remissions currently requires invasive endoscopic evaluation with biopsies, thereby limiting the number of follow-up measurements after therapy initiation and restricting molecular exploration of the mucosa to a set of infracentimetric biopsies. In this context, a noninvasive method of monitoring therapeutic response would be of great benefit to patients. We investigated whether MPIO-αMAdCAM-1 enhanced imaging can detect the effects on MAdCAM-1 expression of statin and antibodies against tumor necrosis factor (TNF) that have been shown to reduce leucocyte infiltration in acute and chronic DSS-induced colitis respectively(*20–22*). First in the acute colitis model, we randomized the mice to receive either atorvastatin *per os* (20 mg/kg) or vehicle (water) for 72 hours after the five days of 2% DSS treatment (**Figure 4A**). Morphological and molecular MRI was performed both before treatment with atorvastatin or vehicle (baseline) and 72 hours thereafter (+72h). Our results demonstrate that atorvastatin treatment reduces MPIO-αMAdCAM-1 induced signal voids at 72 hours (**Figure 4B-D**). Moreover, between baseline and +72h endpoints, MPIO-αMAdCAM-1 induced signal voids diminished in atorvastatin treated mice whereas it slightly increased in vehicle treated mice (**Figure 4E**).

**Figure 4.**
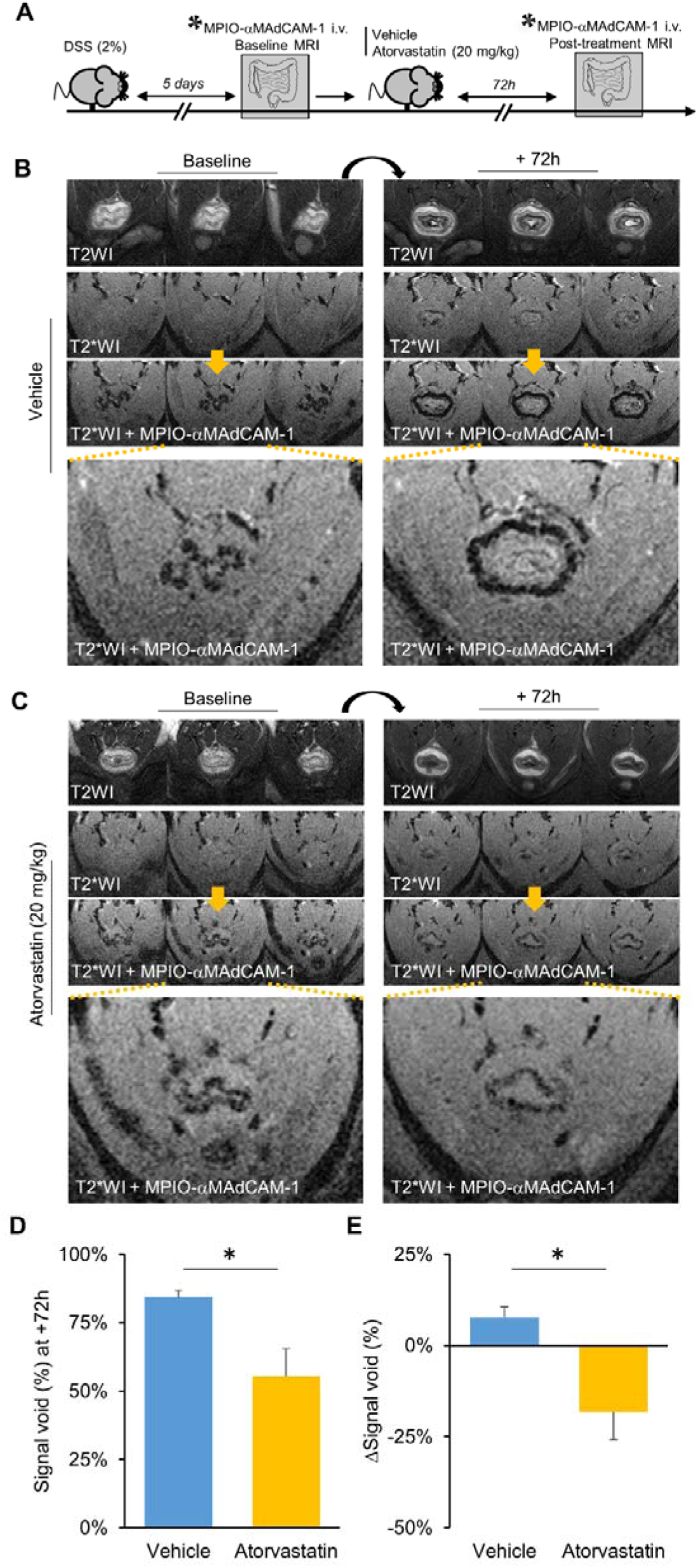
MPIO-αMAdCAM-1 enhanced imaging of the therapeutic response to atorvastatin in the DSS model of acute colitis. (A) Schematic representation of the experiment. (B) Representative T2-weighted, T2*-weighted and T2*-weighted after injection of MPIO-αMAdCAM-1 images in vehicle treated mice before (left) and after 72 hours of vehicle administration (right). (C) Representative T2-weighted, T2*-weighted and T2*-weighted after injection of MPIO-αMAdCAM-1 images in atorvastatin treated mice before (left) and after 72 hours of atorvastatin administration (right). (D) Corresponding quantification of signal void at 72 hours (n=5 for vehicle group and n=4 for atorvastatin group). (E) Corresponding quantification of signal void evolution between baseline and 72 hours (n=5 for vehicle group and n=4 for atorvastatin group). *p<0.05 vs the indicated group.

Then, in the chronic colitis model, we randomized mice to receive either a monoclonal rat anti-mouse anti-TNF antibody (2.5 mg/kg as a single intraperitoneal injection) or an equivalent amount of a control isotype antibody. Morphological and molecular MRI was performed both before treatment (baseline) and 5 days thereafter. In line with the previously reported beneficial effect of anti-TNF treatment in this model, MPIO-αMAdCAM-1 induced signal voids were strongly reduced 5 days after treatment compared to control mice (**Figure 5A-D**). Moreover, MPIO-αMAdCAM-1 induced signal voids more strongly diminished in anti-TNF treated mice compared to control mice (**Figure 5E**). Noteworthy, in both the acute and chronic models of colitis, morphological imaging failed to identify significant changes associated with atorvastatin or anti-TNF treatment, suggesting that MPIO-αMAdCAM-1 enhanced imaging is more sensitive to detect therapeutic response in colitis.

**Figure 5.**
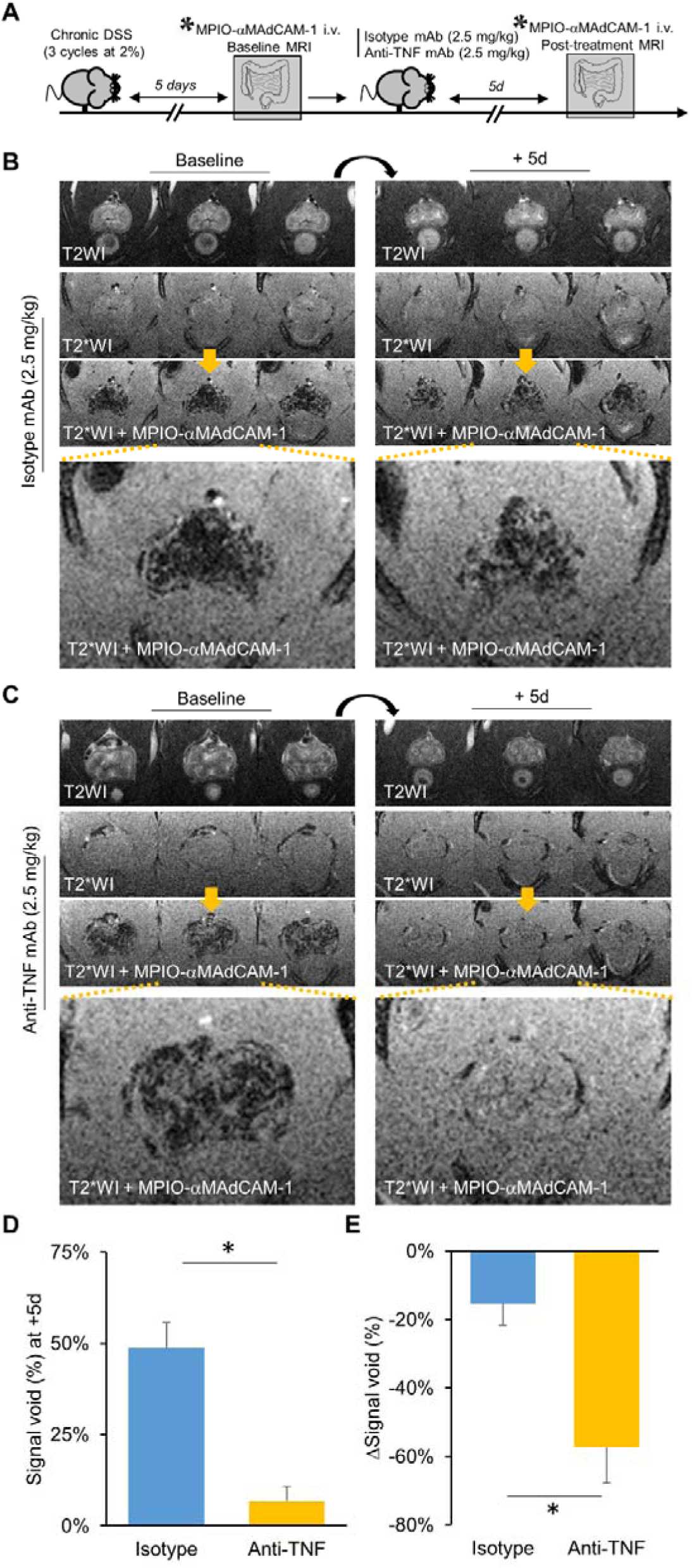
MPIO-αMAdCAM-1 enhanced imaging of the therapeutic response to anti-TNF antibodies in the DSS model of chronic colitis. (A) Schematic representation of the experimental procedure. (B) Representative T2-weighted, T2*-weighted and T2*-weighted after injection of MPIO-αMAdCAM-1 images in control antibody treated mice before (left) and 5 days after control antibody administration (right). (C) Representative T2-weighted, T2*-weighted and T2*-weighted after injection of MPIO-αMAdCAM-1 images in anti-TNF antibody treated mice before (left) and 5 days after anti-TNF antibody administration (right). (D) Corresponding quantification of signal void at 5 days (n=5 per group). (E) Corresponding quantification of signal void evolution between baseline and 5 days (n=5 per group). *p<0.05 vs the indicated group.

### MPIO-**α**MAdCAM-1 allows longitudinal evaluation of mucosal inflammation in the TNBS model of acute colitis

Another interesting advantage of non-invasive imaging is to allow multiple longitudinal evaluations of the same individuals in order to follow more closely the activity of the disease. We wanted to illustrate this possibility by performing longitudinal imaging of mice in another model of acute colitis in mice, induced by intra-rectal administration of TNBS (100 mg/kg in 50% ethanol)(*23*). TNBS couples with high molecular weight proteins in the colon and elicits an acute Th1 driven immunological responses. In this model, we performed axial imaging of the descending colon including T2-weighted sequences and T2*-weighted sequences both before and after MPIO-αMAdCAM-1 injection at different time points in the same animals (at baseline and 2, 4, 7, 10 and 14 days after TNBS exposure, **Figure 6A**). Representative images at 4 days post-TNBS are presented in **Figure 6B**. At this time-point, T2-weighted imaging confirmed acute colitis by revealing a thickening of the colonic mucosa as well as a significant submucosal edema. Post-MPIO-αMAdCAM-1 T2*-weighted sequences shows intense binding in the mucosa in line with active inflammation. Immunohistological analyses in a subset of mice confirmed the binding of numerous MPIO-αMAdCAM-1 in the vessels of the lamina propria of TNBS treated mice, without evidence for extravasated particles (**Figure 6C**). Longitudinal morphological MRI revealed significant mucosal thickening during the first 7 days and submucosal edema during the first 4 days after TNBS exposure (**Figure 6D**). Interestingly, molecular imaging using MPIO-αMAdCAM-1 revealed activated endothelial cells in the colonic mucosa from 2 days to 14 days after TNBS exposure with a peak at 4 days (**Figure 6E**). This highlights the higher sensitivity of molecular imaging to reveal inflammatory changes compared to morphological imaging, especially at 7 and 10 days post-TNBS exposure. These results are summarized in **Figure 6F**.

**Figure 6.**
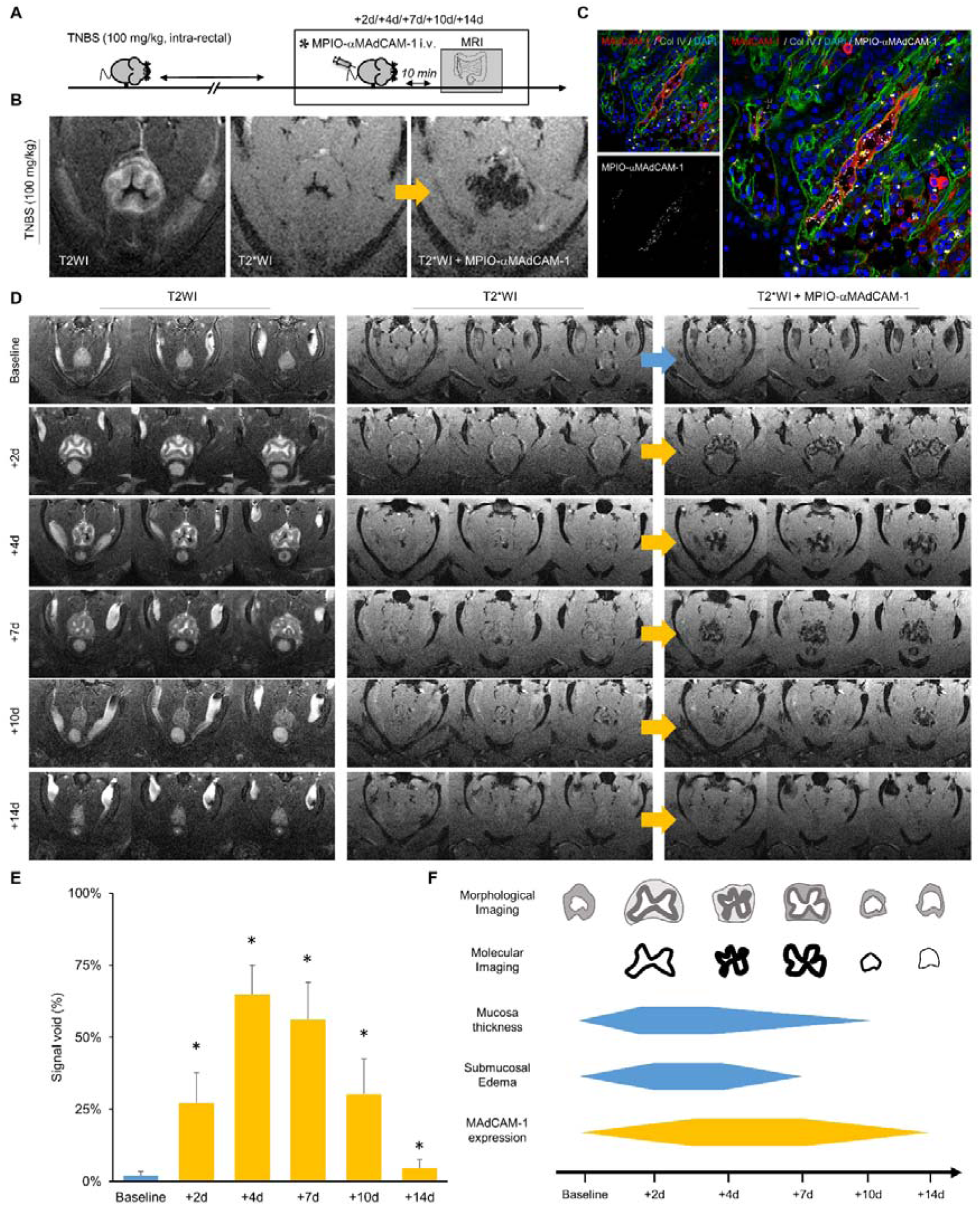
MPIO-αMAdCAM-1 allows longitudinal evaluation of mucosal inflammation in the TNBS model of acute colitis. (A) Schematic representation of the experimental procedure. Mucosal inflammation was evaluated by MRI at baseline, day 2, 4, 7, 10 and 14 after TNBS treatment. (B) Representative T2-weighted, T2*-weighted and T2*-weighted after injection of MPIO-αMAdCAM-1 images at day 4 after TNBS treatment. (C) Representative immunofluorescence images of MPIO-αMAdCAM-1 on the surface of a MAdCAM-1 positive vessel at day 4 after TNBS treatment. (D) Representative T2-weighted, T2*-weighted and T2*-weighted after injection of MPIO-αMAdCAM-1 images in the TNBS model at 6 different time points. (E) Corresponding quantification of signal void (n=8 per group). (F) Summary of the findings in the TNBS model.

## DISCUSSION

We report a new MRI contrast agent allowing non-invasive detection of mucosal inflammation in an ultrasensitive manner (**Figure 7**). The technique presented herein, if successfully applied in humans, holds potential for early diagnosis, longitudinal follow-up and therapeutic response monitoring in diseases involving mucosal tissue, such as IBD. In clinical settings, morphological MRI coupled to molecular imaging of MAdCAM-1 could allow fast, sensitive and non-invasive evaluation of mucosal lesions, thereby alleviating the need for endoscopy and improving patient comfort and safety. The ability to detect MAdCAM-1 could also be key for the identification of patients candidate for MAdCAM-1 targeted therapies such as vedolizumab(*24*). Beyond mucosal lesions, this method has also the potential to improve the diagnosis of other disorders involving dysregulation of MAdCAM-1 such as pancreatic cancers(*25*) or chronic inflammatory liver disease(*26*). The main limitation is the lack of direct clinical translatability of the particles and antibodies that were used in the present study. However, biodegradable MPIOs are now available(*27*) and anti-human MAdCAM-1 antibodies are currently under clinical investigation(*28*), opening the way for clinical translation of our imaging method by coupling the two constructs.

**Figure 7.**
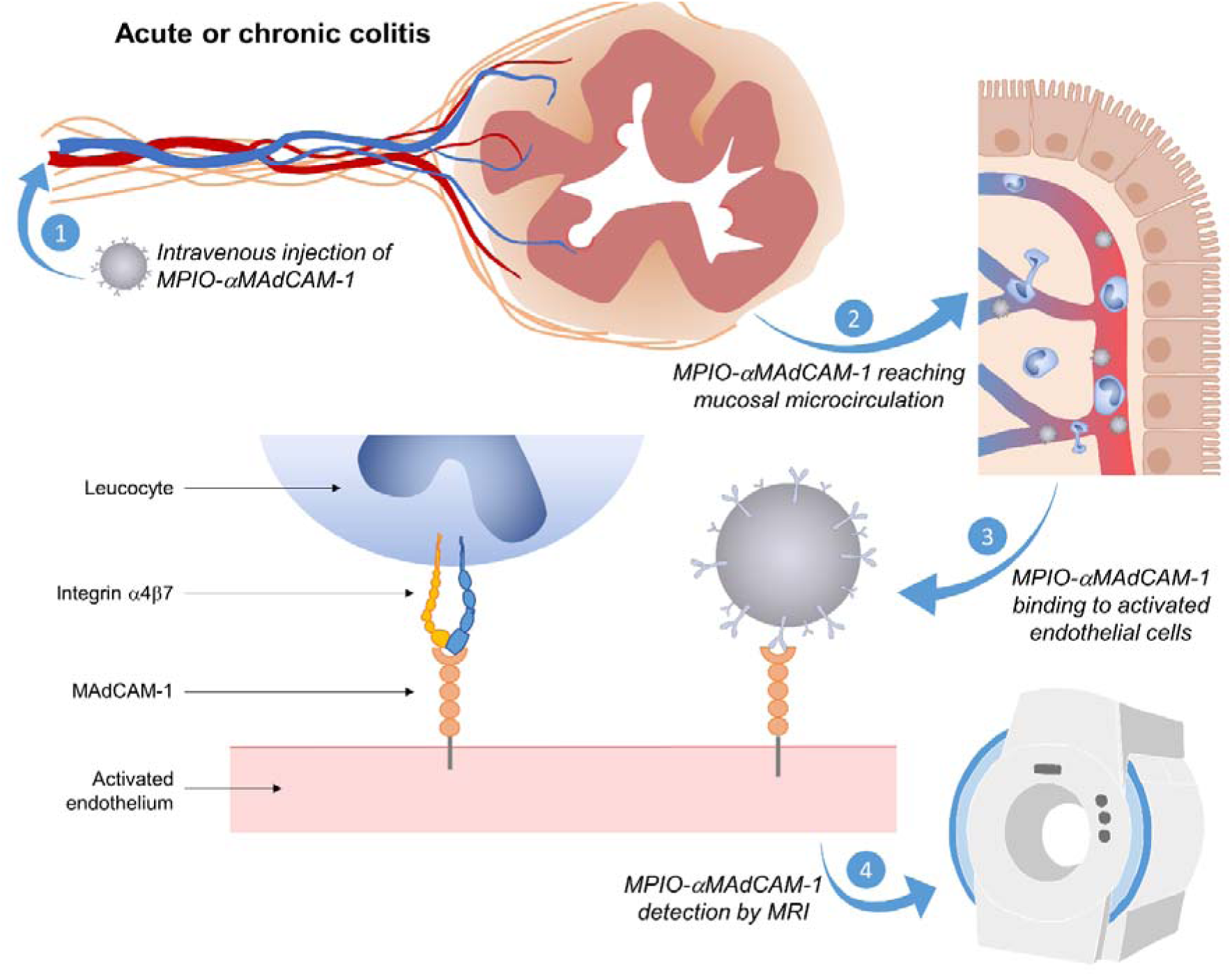
Schematic representation of the main findings.

Since MRI is a widely available technique that is routinely performed in patients affected by mucosal disorders and since our contrast agent can be detected using standard sequences and magnets, the potential impact in patient care could be significant. Only a minor modification of currently performed imaging protocols would be necessary to include a T2*-weighted sequence and therefore to allow MPIO-αMAdCAM-1 detection. Notably, in the context of mucosal inflammation, MRI compares favorably to the other non-invasive methods available for molecular imaging such as positron-emission tomography(*29*) and ultrasounds. A few targeted contrast agents for ultrasounds have been developed (including microbubbles targeted to adhesion molecules)(*30, 31*), but there are several drawbacks that disqualify ultrasound as a universally suitable method for detecting mucosal inflammation, such as an exploration limited to superficial tissues and the frequent presence of obstacles (such as gas) that precludes evaluation of some organs or intestinal segments. Combining ultrasounds with laser light stimulation, photoacoustic imaging allows imaging of mucosal inflammation in Crohn’s disease without exogenous contrast but remains impaired by the same limitations than ultrasounds(*32*). Positron-emission tomography, using 18F-Fluorodeoxyglucose or immuno-PET techniques, is theoretically suitable for imaging of mucosal tissue in the whole body but is limited by its low spatial resolution, its long acquisition time and its use of ionizing radiation, that precludes its repeated use in patients with chronic diseases(*29*). MRI is devoid of these drawbacks and the use of MPIO as contrast-carrying particles compensates for its low sensitivity(*33*), thus allowing sensitive imaging of mucosal inflammation.

In summary, this experimental study demonstrates that non-invasive and high-spatial-resolution mapping of mucosal inflammation is feasible *in vivo* using MPIO-αMAdCAM-1 enhanced MRI, with potential applications in early diagnosis, longitudinal follow-up and therapeutic response monitoring in diseases involving mucosal tissues.

## MATERIAL AND METHODS

### Animals

All experiments were performed on 8-week-old male Swiss mice (Janvier, France). Animals were maintained under specific pathogen-free conditions at the Centre Universitaire de Ressources Biologiques (CURB, Basse-Normandie, France) and all had free access to food and tap water. A 6-hour fasting period was observed before MRI procedures. Experiments were approved by the local ethical committee (n° 2889, « Imagerie par résonnance magnétique de l’inflammation intestinale ») and were performed in accordance with the French (Decree 87/848) and the European Communities Council (Directive 2010/63/EU) guidelines.

### Targeting-moiety conjugation to MPIO

Microparticles of iron oxide (MPIO; diameter 1.08 μm) with p-toluenesulphonyl reactive surface groups (Invitrogen) were covalently conjugated to purified monoclonal rat anti-mouse monoclonal antibodies for MAdCAM-1. Briefly, purified monoclonal rat anti-mouse antibodies for MAdCAM-1 (clone MECA-367; BD-Biosciences) or control isotype (clone R35-95; BD-Biosciences) were covalently conjugated to MPIO in borate buffer (pH 9.5), by incubation at 37 °C for 48 h. 40 μg of targeting molecule was used for the coating of 1 mg of reactive MPIO. MPIO were then washed in phosphate buffered saline (PBS) containing 0.5% bovine serum albumin (BSA) at 4 °C and incubated for 24 h at room temperature, to block the remaining active groups. MPIOs were rinsed in PBS (0.1% BSA). To disperse MPIO aggregates, a sonication procedure at low intensity was performed one time, immediately after antibody labelling, for 60 seconds. Thereafter, conjugated MPIO were stored at 4°C in a PBS buffer under constant agitation to prevent settling and aggregate formation. We used each MPIO batches up to 1 month after antibody labeling (8 batches were produced for this study). Finally, monodisperse MPIO (as assessed by confocal microscopy) were stored at 4°C. Labelling efficacy was characterized by flow cytometry and confocal microscopy. Using a similar protocol of antibody conjugation, Jefferson et al. measured that the density of targeting antibodies on the surface of MPIO was 27,100 ± 1920 molecules per MPIO, equivalent to 8500 molecules μm. (*34*)

### MPIO injection

Anesthesia was induced using 5% isoflurane (Aerrane, Baxter) and maintained using 2% isoflurane in a mixture of O2/N2O (30%/70%). A tail vein catheter was placed in one of the lateral veins to allow intravenous injection. Mice received intravenous injection of 1.0 mg/kg (equivalent Fe) of conjugated MPIO for contrast-enhanced MRI through the tail vein catheter. Imaging was performed 10 minutes after MPIO administration.

### Magnetic Resonance Imaging (MRI)

MRI Experiments were carried out on a Pharmascan 7 T/12 cm system using surface coils (Bruker, Germany). Anesthesia was induced using 5% isoflurane (Aerrane, Baxter) and maintained using 2% isoflurane in a mixture of O2/N2O (30%/70%) to ensure immobility during MRI. Abdominal imaging was performed using a T2*-weighted 2D sequence (Fast low angle shot) with respiratory gating and the following parameters: TE/TR 5.4 ms/variable, flip angle 25°, with a matrix of 256 × 256, a field of view of 20 × 20 mm leading to a 78*78 µm spatial resolution and a slice thickness of 0.5 mm. Both axial and sagittal abdominal imaging were performed with the same sequence parameters. Molecular imaging of the colon was performed using a T2*-weighted 2D sequence (Fast low angle shot) without gating and using the following parameters: TE/TR 5.4 ms/300 ms, flip angle 35°, with 4 excitations, a matrix of 300 × 300, a field of view of 20 × 20 mm leading to a 67*67 µm spatial resolution and a slice thickness of 0.35 mm (acquisition time: 6 min). Morphological imaging of the colon was performed using a T2-weighted 2D sequence (Rapid Acquisition with Relaxation Enhancement) without gating and using the following parameters: TE/TR 36 ms/2500 ms, flip angle 90°, refocusing pulse 180°, acceleration factor 8 with 3 excitations, a matrix of 300 × 300, a field of view of 20 × 20 mm leading to a 67*67 µm spatial resolution and a slice thickness of 0.75 mm (acquisition time: 4 min 37 s). Although all MPIO injections were performed outside the magnet (to avoid sedimentation of the MPIO due to the magnetic field of the MRI), the positions of the mice during pre-MPIO and post-MPIO acquisitions did not change (the mice were placed exactly at the same position and were not removed from the animal holder).

### MRI Analysis

All analyses were performed using ImageJ v1.52 (National institute of health) in a similar manner as previously described for the brain, heart and kidney(*14*). First, the mucosa was segmented using either the pre-contrast T2*-weighted images (for abdominal imaging in LPS or saline treated mice) or the anatomical T2-weighted images for the colon (otherwise). Then a manual threshold is applied using ImageJ software and the signal void percentage is computed by dividing the volume of voxels from the mucosa with a signal inferior to the threshold by the volume of voxels with a signal superior to the threshold.

### Competition experiments

When appropriate, mice received 2.5 mg/kg of anti-MAdCAM-1 antibody or control isotype IgG 15 minutes before MPIO-αMAdCAM-1 injection to block the MPIO-binding sites. MRI was thereafter performed as described below.

### LPS-induced endothelial activation

Mice received an intra-peritoneal (i.p.) injection of LPS from E. coli (2.5 mg/kg, 0111:B4, Sigma-Aldrich, Lisle d’Abeau, France). MRI was performed 48 hours after LPS administration.

### DSS model of acute colitis

DSS-treated mice received 1% to 4% (weight/volume) DSS (40,000 MW, Alfa Aesar, Thermo Fisher, France) in drinking water for 5 days to induce an acute episode of colitis. Drink and food were provided ad libitum. DSS, food consumption, and body weight were measured daily. Mice were evaluated using a disease activity index based on weight loss (0: none, 1: 1-5%, 2: >5-10%, 3: >10-20%, 4: >20%), stool consistency (0: well-formed pellets, 2: loose, pasty stools, 4: diarrhea) and rectal bleeding (0: none, 2: mild bleeding, 4: gross bleeding). Presented data are the sum of the three scores. Healthy controls received tap water. When appropriate, mice received atorvastatin (20 mg/kg *per os*, Tahor, Pfizer) for 3 days and control mice received the same volume of water *per os*.

### DSS model of chronic colitis

DSS-treated mice received 3 cycles of 2% (weight/volume) DSS (40,000 MW, Alfa Aesar, Thermo Fisher, France) in drinking water for 5 days. Each cycle was separated by 7 days without DSS in the drinking water. When appropriate, mice received an injection of rat anti-mouse tumor necrosis factor (TNF) antibody (2.5 mg/kg intraperitoneal, clone MP6-XT22, Thermo Fisher) or a control isotype antibody (IgG1 kappa, Thermo Fisher).

### TNBS model of acute colitis

Mice were anesthetized using 5% isoflurane and a canula was inserted through the anus into the colon for 4 cm, up to the tip of a 4*0.4 cm canula (Ecimed, Boissy-Saint-Léger, France). Then 100 µl of a 2.5% solution (weight/volume) of TNBS (Sigma-Aldrich, Lisle d’Abeau, France) reconstituted in 50% ethanol were slowly injected. Mice were kept head down for 60 seconds to avoid reflow of the solution and returned to their cage.

### Immunohistochemistry

Deeply anesthetized mice were transcardially perfused with cold heparinized saline (15 mL) followed by 150 mL of fixative (PBS 0.1 M. pH 7.4 containing 2% paraformaldehyde and 0.2% picric acid). Ileum or colons were post-fixed (18 hours; 4°C) and cryoprotected (sucrose 20% in veronal buffer; 24 hours; 4°C) before freezing in Tissue-Tek (Miles Scientific, Naperville, IL, USA). Cryomicrotome-cut transversal sections (10 µm) were collected on poly-lysine slides and stored at −80°C before processing. Sections were co-incubated overnight with rat monoclonal anti-mouse MAdCAM-1 (1:1000; clone MECA-367 from BD Biosciences) and goat anti-collagen-type IV (1:1500; Southern Biotech) in veronal buffer (pH 7.4). Primary antibodies were revealed using Fab’2 fragments of Donkey anti-rat or goat IgG linked to FITC or TRITC (1:500, Jackson ImmunoResearch, West Grove, USA). Washed sections were coverslipped with antifade medium containing DAPI and images were digitally captured using a Leica DM6000 microscope-coupled coolsnap camera and visualized with Metavue 5.0 software (Molecular Devices, USA) and further processed using ImageJ 1.52 software. Quantitative analyses were performed using three randomly selected slice per mice. A vessel was considered MAdCAM-1 positive when the fluorescence signal was distinguishable from the background. All analyses were performed at 200 × magnification and blinded to the experimental data.

### Flow cytometry

Uncoated MPIO and MPIO-αMAdCAM-1 were analyzed on a FACSVerse (Becton Dickinson) equipped with its accompanying software (FACSuite, Becton Dickinson) on 10,000 events after gating by FSC and SSC the MPIO population. When stated, anti-rat Fab’2 antibodies conjugated with phycoerythrin (PE) were incubated with the MPIO for 30 min and washed before running the samples through the cytometer.

### Confocal analysis of MPIO conjugation

Laser-scanning confocal microscopy was performed using an inverted Leica SP5 confocal microscope (Leica Microsystems SAS) equipped with an Argon Gas laser and a X40 NA=1.4 oil immersion objective. For tetramethylrhodamine (TRITC) detection, excitation was set at 561 nm and emission filters between 575 and 625 nm. Field of view was set at 1 µm × 1 µm with a 1024×1024 planar matrix (pixel size= 97.6 nm × 97.6 nm). Analyses of the images were performed using imageJ 1.52 software. When stated, anti-rat Fab’2 antibodies conjugated with TRITC were incubated with the MPIO for 30 min and then washed before imaging. MPIO fluorescence was evaluated on all visible particles from 3 randomly selected slice, leading to sample sizes between 180 and 220 particles.

### Statistical analysis

Results are presented as the mean ± SD. Statistical analyses were performed using Mann-Whitney’s U-test. When more than two groups were compared, statistical analyses were performed using Kruskal-Wallis (for multiple comparisons) followed by Mann-Whitney’s U-test. When comparing two groups, a p-value < 0.05 was considered to be significant. For longitudinal analyses, statistical analyses were performed using Wilcoxon signed-rank test with baseline as the first value and without correction for multiple testing.

## Supplementary Materials

**Figure S1.**
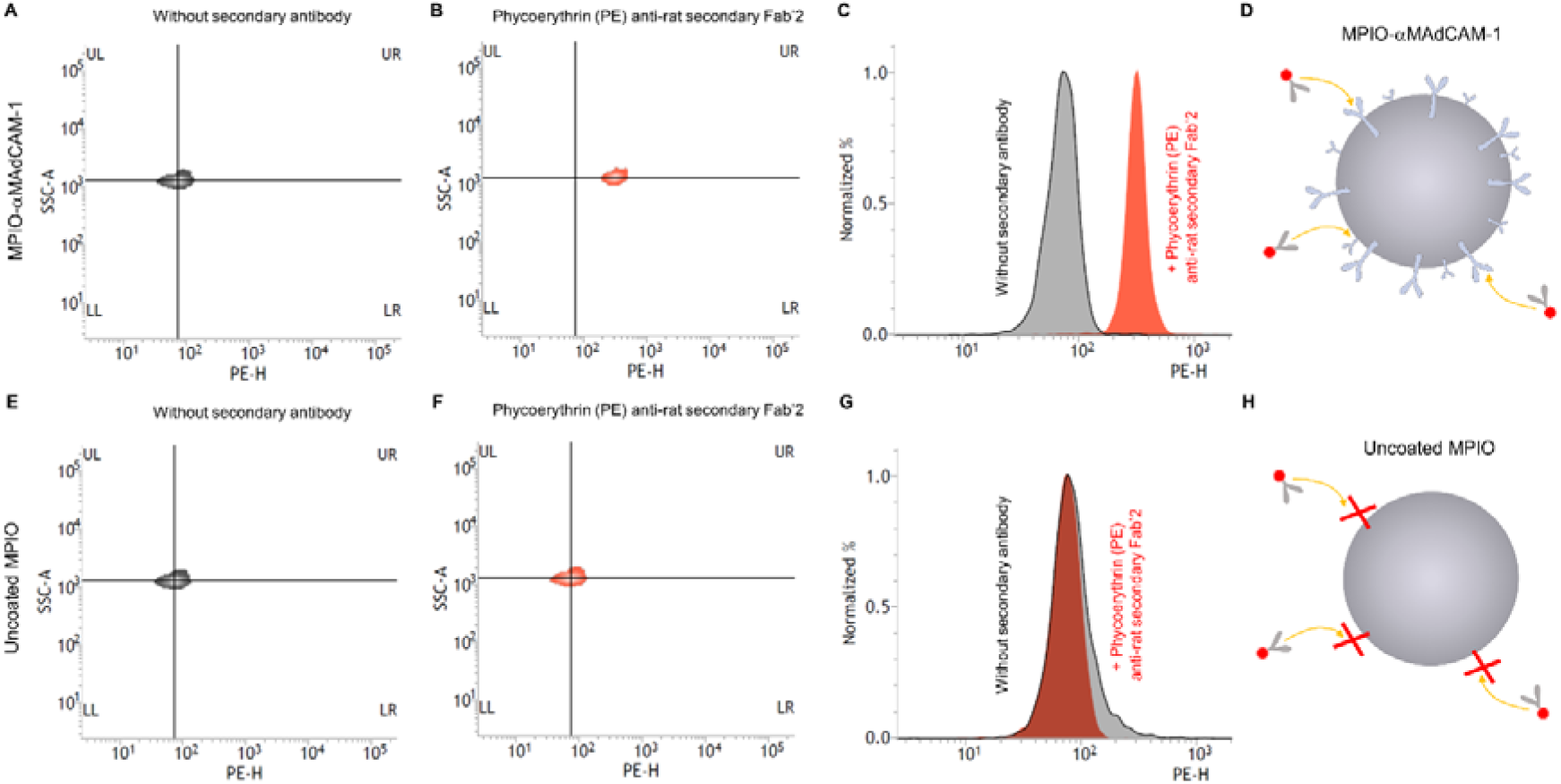
Effective coating of MPIO with anti-MAdCAM-1 monoclonal antibodies as assessed by flow cytometry. 2D density plots of MPIO-αMAdCAM-1 (A) without and (B) with secondary anti-rat Fab’2 antibodies. (C) Histogram analysis of the data presented in A and B showing a shift in the fluorescence of the MPIO-αMAdCAM-1 population in presence of the secondary anti-rat Fab’2, corresponding to the binding of the secondary antibodies on the MPIO-αMAdCAM-1. (D) Schematic representation of the results of A, B and C. (E), (F), (G) and (H) are the same as A, B, C and D but using uncoated MPIO. The secondary anti-rat Fab’2 antibodies do not bind uncoated MPIO, explaining the absence of a shift in the fluorescence when the MPIO were incubated with the secondary antibodies.

**Figure S2.**
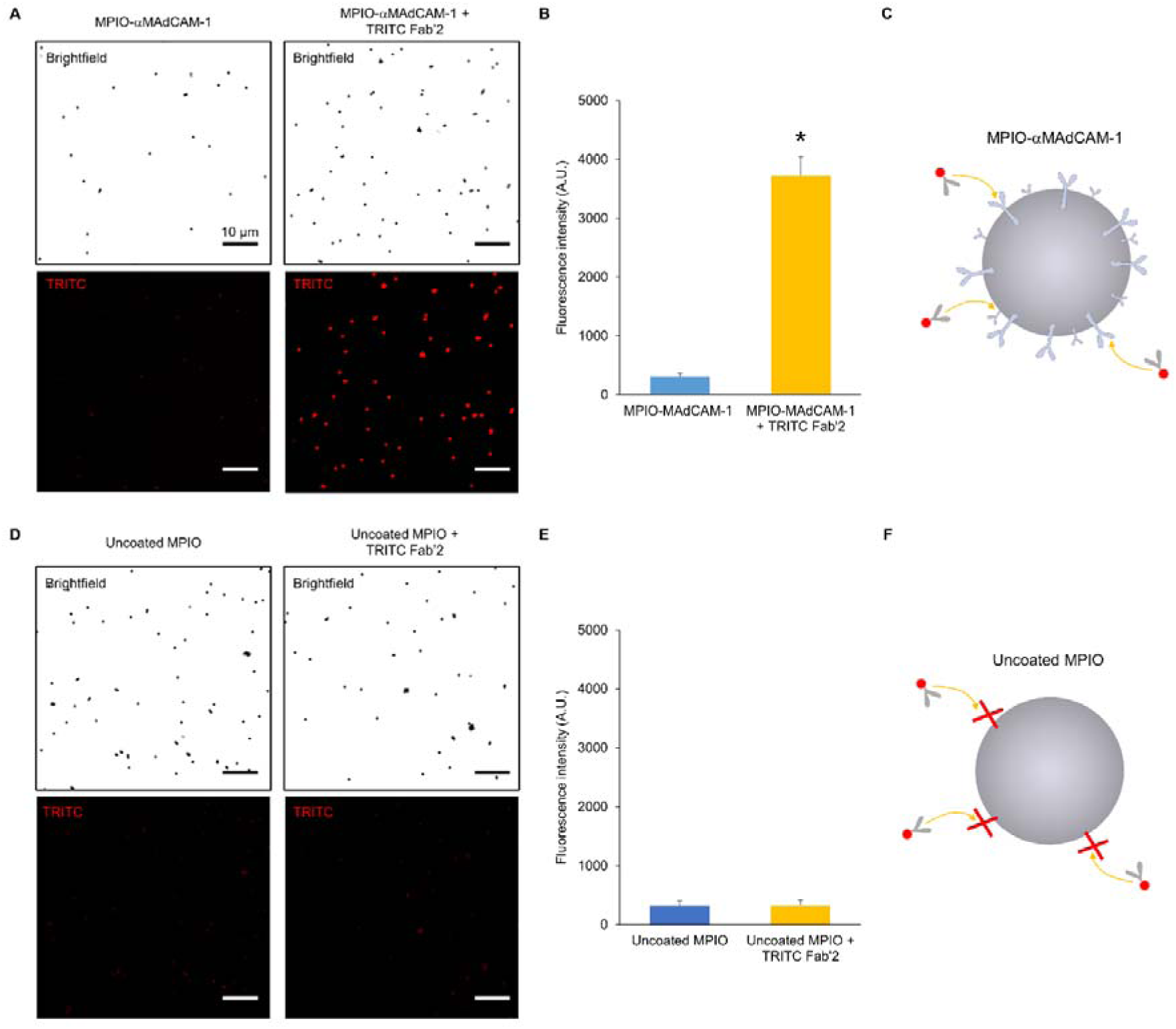
Effective coating of MPIO with anti-MAdCAM-1 monoclonal antibodies as assessed by confocal microscopy. (A) Representative confocal microscopy images (top: brightfield by reflectance imaging; bottom: tetramethylrhodamine (TRITC) fluorescence, red) of MPIO-αMAdCAM-1 without (left) or with (right) TRITC anti-rat secondary Fab’2. (B) Corresponding analyses (n=∼200 per group) showing a higher fluorescence of the MPIO-αMAdCAM-1 population in presence of the secondary anti-rat Fab’2, corresponding to the binding of the secondary antibodies on the MPIO-αMAdCAM-1. (C) Schematic representation of the results of A and B. (D), (E), (F) are the same as A, B and C but using uncoated MPIO. The secondary TRITC anti-rat Fab’2 antibodies do not bind uncoated MPIO, explaining the similar fluorescence between the uncoated MPIO that were incubated with the secondary antibodies and the uncoated MPIO that were not, as shown in (E) (n=∼200 per group).*p<0.05.

**Figure S3.**
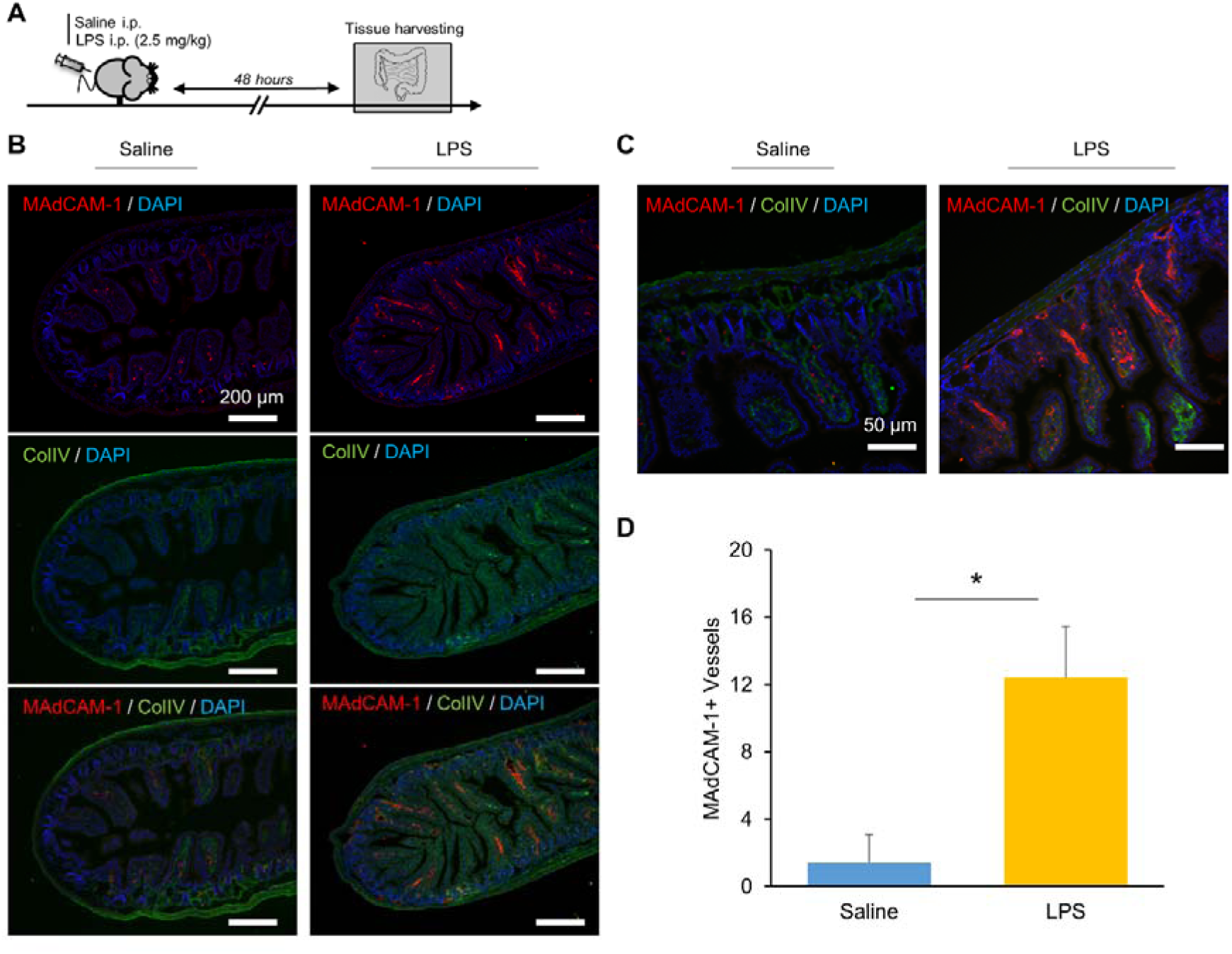
Intraperitoneal LPS administration induces MAdCAM-1 overexpression in the ileum. (A) Schematic representation of the experimental procedure. (B) Representative immunofluorescence images of MAdCAM-1 (red), collagen type IV (green) and DAPI (blue) in the ileum of mice 48 hours after LPS administration. (C) Representative high magnification image that was used to count the number of MAdCAM-1 expressing vessel. (D) Corresponding quantification. *p<0.05 versus saline.

**Figure S4.**
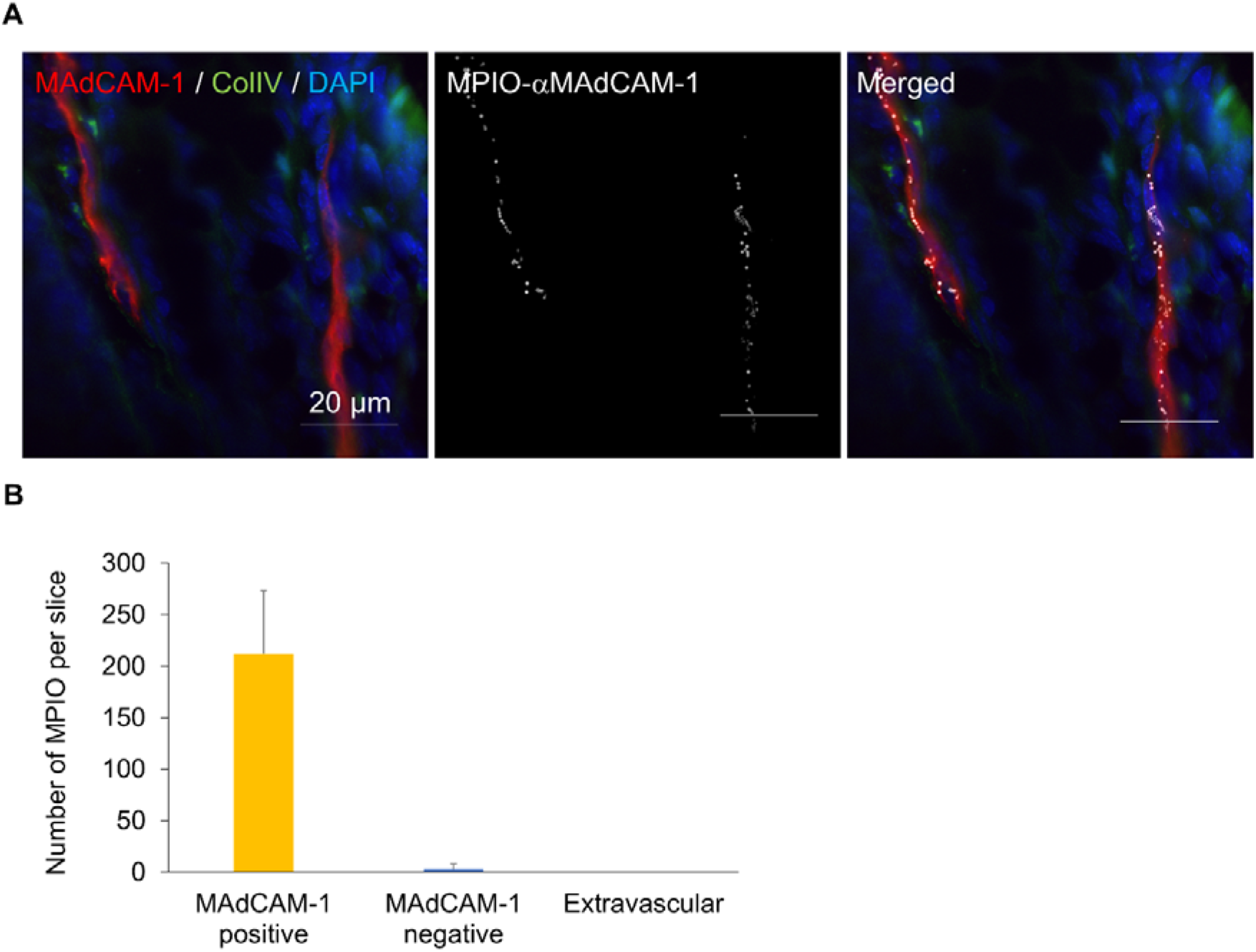
MPIO-αMAdCAM-1 bind to MAdCAM-1 expressing vessels after intravenous injection in the LPS model. (A) Representative immunofluorescence image of MAdCAM-1 (red), collagen type IV (green), DAPI (blue) and MPIO-αMAdCAM-1 10 minutes after intravenous injection in the ileum of a mouse 48 hours after intraperitoneal LPS injection (2.5 mg/kg). (B) Corresponding quantification of MPIO-αMAdCAM-1 according to its localization (n=4).

**Figure S5.**
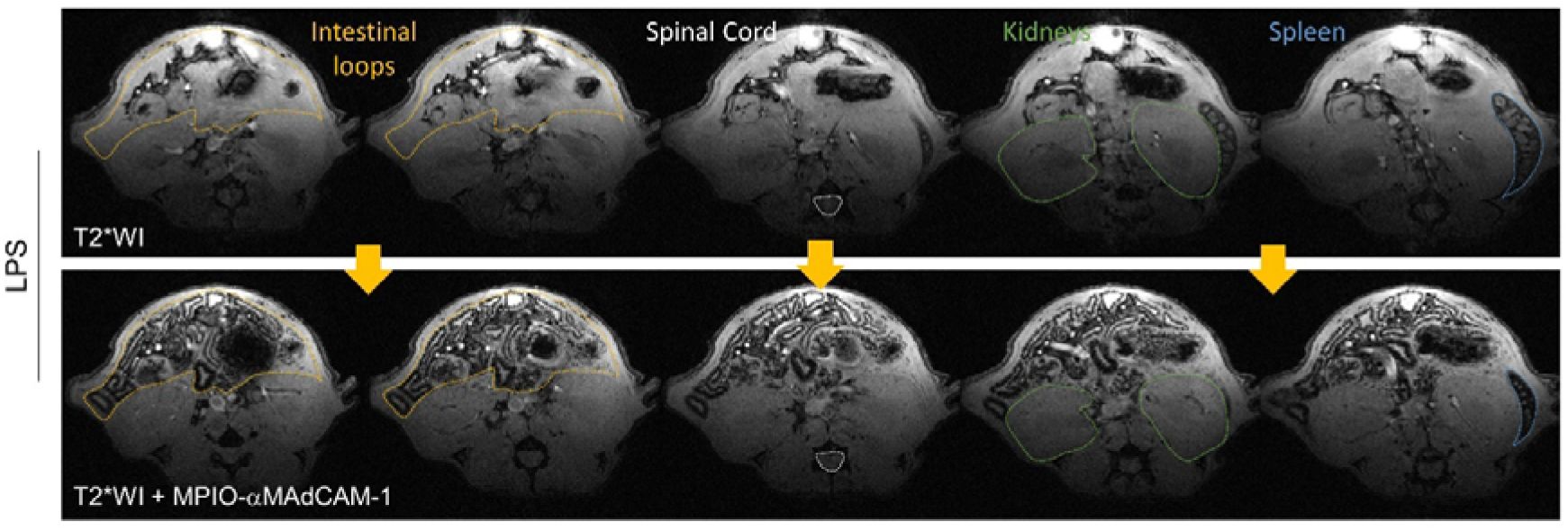
Multislice T2*-weighted images before and after MPIO-αMAdCAM-1 administration in an LPS treated mouse. Relevant anatomical structures are labeled.

**Figure S6.**
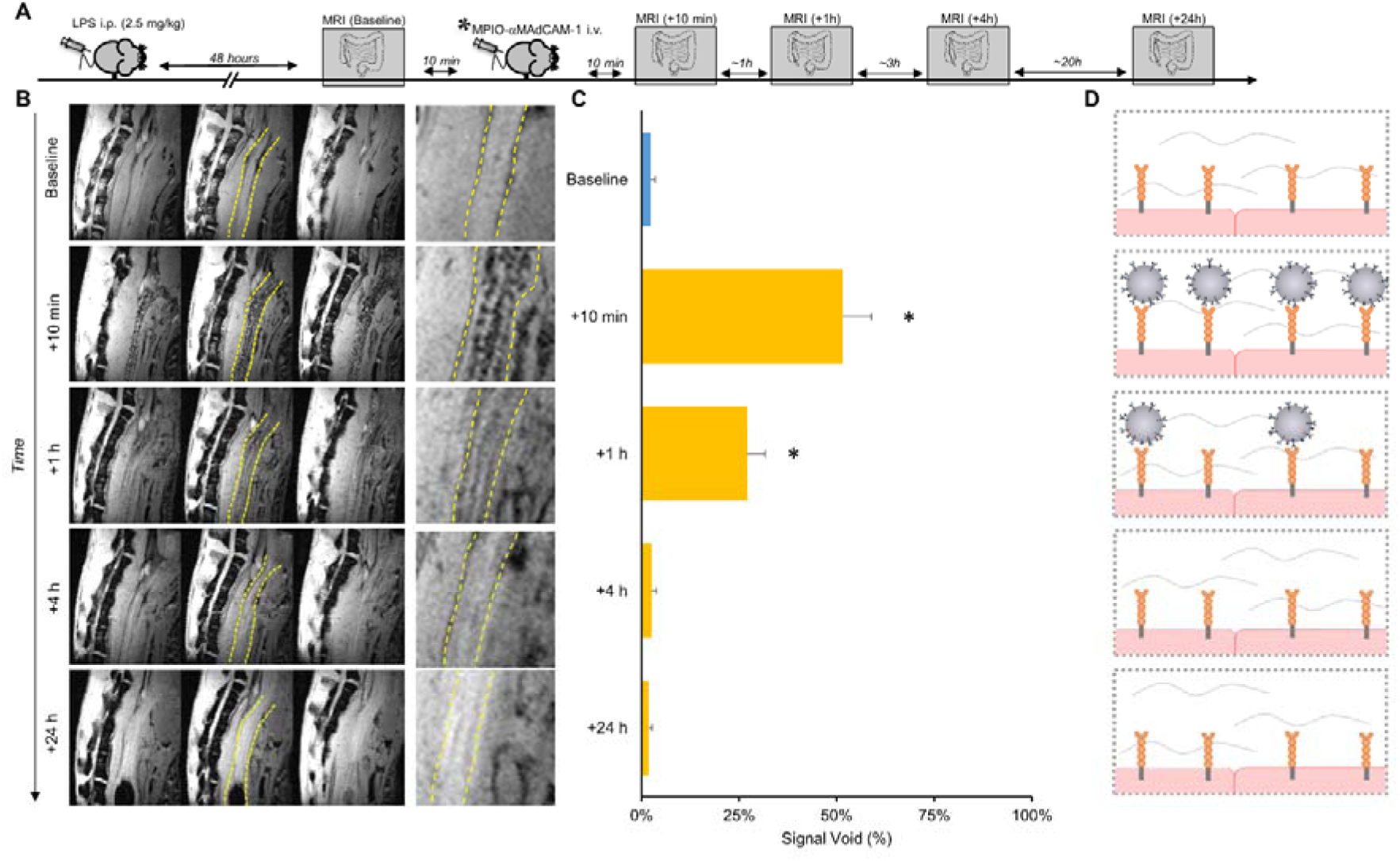
Longitudinal imaging after a single injection of MPIO-αMAdCAM-1 in LPS treated mice. (A) Schematic representation of the experimental procedure. (B) Representative T2*-weighted images both before and after injection of MPIO-αMAdCAM-1 at different time-points (baseline, 10 min, 1h, 4h and 24h post-MPIO-αMAdCAM-1 injection). Images are centered on the descending colon (sagittal slices) which is a fixed intestinal loop, thereby easing longitudinal quantification. (C) Corresponding quantification. (D) Schematic representation of the results.

**Figure S7.**
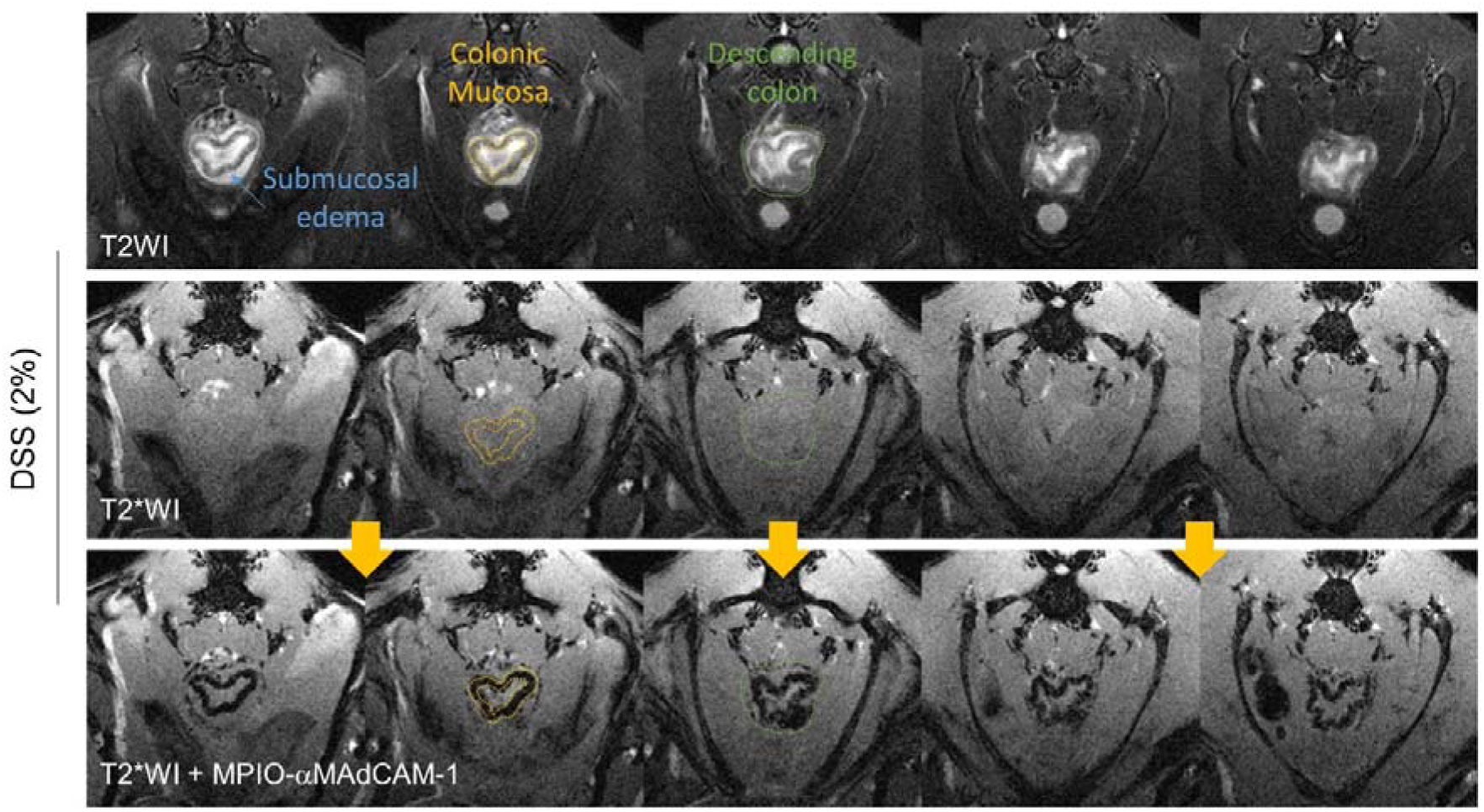
Multislice T2 and T2*-weighted images before and after MPIO-αMAdCAM-1 administration in the DSS model of acute colitis. Relevant anatomical structures are labeled.

**Figure S8.**
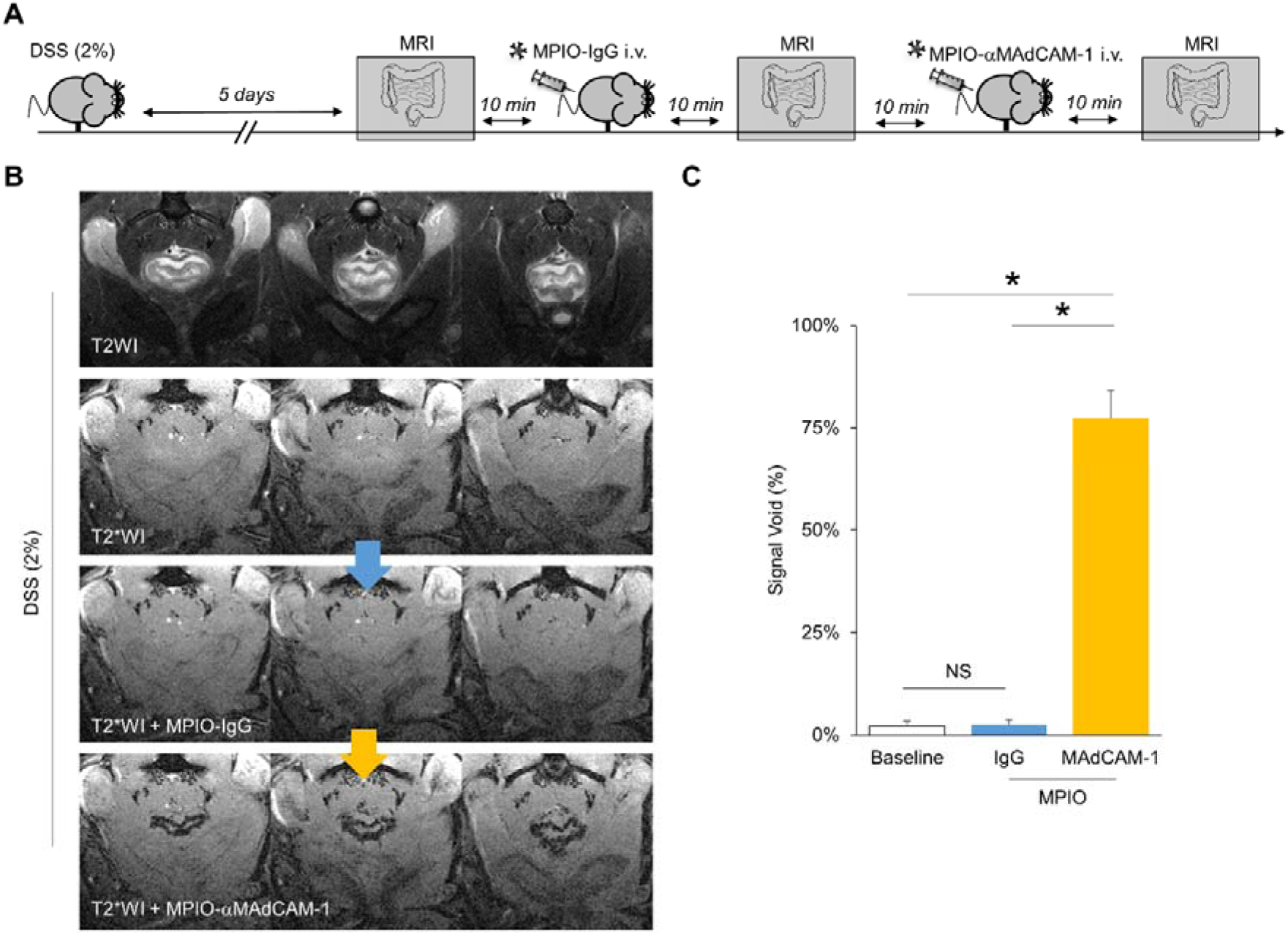
MPIO-αMAdCAM-1 bind specifically to the inflammatory mucosa in the DSS model of acute colitis. (A) Schematic representation of the experimental procedure. (B) Representative T2-weighted, T2*-weighted, T2*-weighted after injection of MPIO-IgG and T2*-weighted after injection of MPIO-αMAdCAM-1 images. (C) Corresponding quantification showing that, unlike MPIO-IgG, injection of MPIO-αMAdCAM-1 induce significant signal voids in the mucosa of mice in the DSS model (n=5 per group). *p<0.05 versus the indicated group.

**Figure S9.**
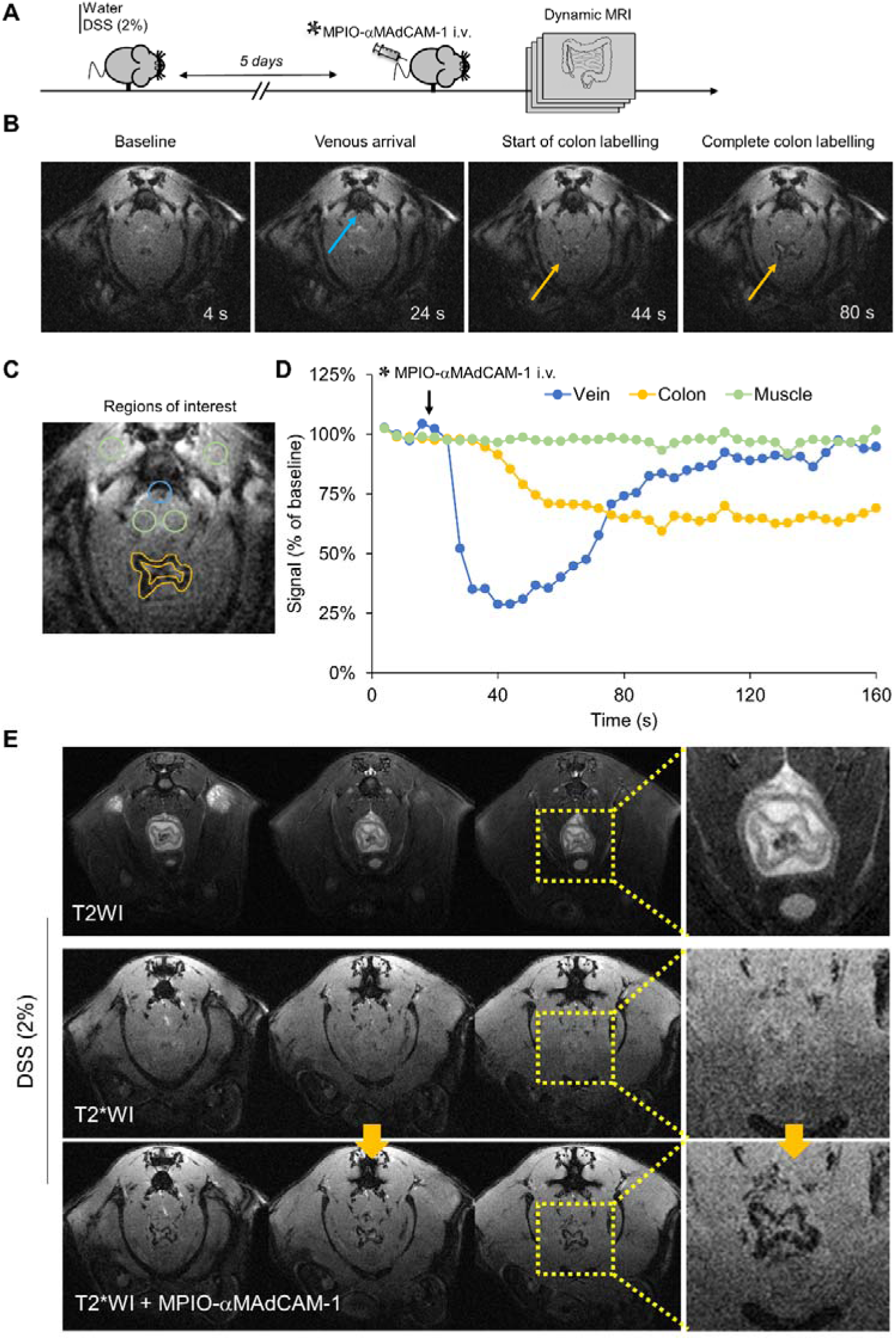
Analysis of the binding kinetic of MPIO-αMAdCAM-1 in the DSS model of acute colitis. (A) Schematic representation of the experimental procedure. Dynamic MRI consisted of 1 image every 4 seconds. (B) Representative T2*-weighted images and key time points before and after intravenous injection of MPIO-αMAdCAM-1. (C) Drawing of the regions of interest that were used for quantification. (D) Evolution of MRI signal over time in the different regions of interest (vein in blue, colon in yellow and muscle in green). (E) Representative T2-weighted, T2*-weighted and T2*-weighted after injection of MPIO-αMAdCAM-1. All the images and quantification presented in this figure are from the same mouse. The results were consistent over the 3 mice performed for this experiment.

## Author contributions

APF, SMDL, AR, AGB, MJW, MFN and MG conceptualized the study. APF, SMDL and MG performed experimental procedures. APF and MG performed statistical analyses. SMDL and MG prepared the figures. MG wrote the original draft. APF, SMDL, AR, AGB, MJW, MFN, DV and FD participated in draft reviewing. DV, FD and MG secured the funding.

## Data and materials availability

All data associated with this study are present in the paper or the Supplementary Materials.

## Supplementary Movie Legends

**Movie S1. MPIO-**α**MAdCAM-1 binds to the activated endothelium of the gastrointestinal tract.** Representative pre and post-contrast T2*-weighted images of the abdomen in a mouse, 48 hours after intraperitoneal LPS administration.

**Movie S2. Binding kinetic of MPIO-**α**MAdCAM-1 in the DSS model of acute colitis.** Representative fast T2*weighted images of the inflamed colon during MPIO-αMAdCAM-1 administration in a mouse after 5 days of 2% DSS treatment.

## REFERENCES

1. H. F. Helander, L. Fandriks, Surface area of the digestive tract - revisited. Scand. J. Gastroenterol. 49, 681–689 (2014).

2. S. C. Ng, H. Y. Shi, N. Hamidi, F. E. Underwood, W. Tang, E. I. Benchimol, R. Panaccione, S. Ghosh, J. C. Y. Wu, F. K. L. Chan, J. J. Y. Sung, G. G. Kaplan, Worldwide incidence and prevalence of inflammatory bowel disease in the 21st century: a systematic review of population-based studies. Lancet 390, 2769–2778 (2018).

3. S. A. Taylor, S. Mallett, G. Bhatnagar, R. Baldwin-Cleland, S. Bloom, A. Gupta, P. J. Hamlin, A. L. Hart, A. Higginson, I. Jacobs, S. McCartney, A. Miles, C. D. Murray, A. A. Plumb, R. C. Pollok, S. Punwani, L. Quinn, M. Rodriguez-Justo, Z. Shabir, A. Slater, D. Tolan, S. Travis, A. Windsor, P. Wylie, I. Zealley, S. Halligan, Diagnostic accuracy of magnetic resonance enterography and small bowel ultrasound for the extent and activity of newly diagnosed and relapsed Crohn’s disease (METRIC): a multicentre trial. The lancet. Gastroenterology & hepatology 3, 548–558 (2018).

4. A. Miles, G. Bhatnagar, S. Halligan, A. Gupta, D. Tolan, I. Zealley, S. A. Taylor, M. investigators, Magnetic resonance enterography, small bowel ultrasound and colonoscopy to diagnose and stage Crohn’s disease: patient acceptability and perceived burden. Eur. Radiol. 29, 1083–1093 (2019).

5. A. J. Greenup, B. Bressler, G. Rosenfeld, Medical Imaging in Small Bowel Crohn’s Disease-Computer Tomography Enterography, Magnetic Resonance Enterography, and Ultrasound: “Which One Is the Best for What?”. Inflamm. Bowel Dis. 22, 1246–1261 (2016).

6. S. Zundler, E. Becker, L. L. Schulze, M. F. Neurath, Immune cell trafficking and retention in inflammatory bowel disease: mechanistic insights and therapeutic advances. Gut 68, 1688–1700 (2019).

7. M. F. Neurath, Targeting immune cell circuits and trafficking in inflammatory bowel disease. Nat. Immunol. 20, 970–979 (2019).

8. F. Colotta, P. Allavena, A. Sica, C. Garlanda, A. Mantovani, Cancer-related inflammation, the seventh hallmark of cancer: links to genetic instability. Carcinogenesis 30, 1073–1081 (2009).

9. M. Gauberti, A. P. Fournier, F. Docagne, D. Vivien, S. Martinez de Lizarrondo, Molecular Magnetic Resonance Imaging of Endothelial Activation in the Central Nervous System. Theranostics 8, 1195–1212 (2018).

10. A. P. Fournier, A. Quenault, S. Martinez de Lizarrondo, M. Gauberti, G. Defer, D. Vivien, F. Docagne, R. Macrez, Prediction of disease activity in models of multiple sclerosis by molecular magnetic resonance imaging of P-selectin. Proc. Natl. Acad. Sci. U. S. A. 114, 6116–6121 (2017).

11. M. Gauberti, A. Montagne, O. A. Marcos-Contreras, A. Le Behot, E. Maubert, D. Vivien, Ultra-sensitive molecular MRI of vascular cell adhesion molecule-1 reveals a dynamic inflammatory penumbra after strokes. Stroke 44, 1988–1996 (2013).

12. A. Montagne, M. Gauberti, R. Macrez, A. Jullienne, A. Briens, J. S. Raynaud, G. Louin, A. Buisson, B. Haelewyn, F. Docagne, G. Defer, D. Vivien, E. Maubert, Ultra-sensitive molecular MRI of cerebrovascular cell activation enables early detection of chronic central nervous system disorders. Neuroimage 63, 760–770 (2012).

13. S. Wirtz, V. Popp, M. Kindermann, K. Gerlach, B. Weigmann, S. Fichtner-Feigl, M. F. Neurath, Chemically induced mouse models of acute and chronic intestinal inflammation. Nat. Protoc. 12, 1295–1309 (2017).

14. J. Belliere, S. Martinez de Lizarrondo, R. P. Choudhury, A. Quenault, A. Le Behot, C. Delage, D. Chauveau, J. P. Schanstra, J. L. Bascands, D. Vivien, M. Gauberti, Unmasking Silent Endothelial Activation in the Cardiovascular System Using Molecular Magnetic Resonance Imaging. Theranostics 5, 1187–1202 (2015).

15. B. Chassaing, J. D. Aitken, M. Malleshappa, M. Vijay-Kumar, Dextran sulfate sodium (DSS)- induced colitis in mice. Curr. Protoc. Immunol. 104, Unit 15.25. (2014).

16. D. Bettenworth, A. Bokemeyer, M. Baker, R. Mao, C. E. Parker, T. Nguyen, C. Ma, J. Panes, J. Rimola, J. G. Fletcher, V. Jairath, B. G. Feagan, F. Rieder, T. Stenosis, C. Anti-Fibrotic Research, Assessment of Crohn’s disease-associated small bowel strictures and fibrosis on cross-sectional imaging: a systematic review. Gut 68, 1115–1126 (2019).

17. J. F. Colombel, R. Panaccione, P. Bossuyt, M. Lukas, F. Baert, T. Vanasek, A. Danalioglu, G. Novacek, A. Armuzzi, X. Hebuterne, S. Travis, S. Danese, W. Reinisch, W. J. Sandborn, P. Rutgeerts, D. Hommes, S. Schreiber, E. Neimark, B. Huang, Q. Zhou, P. Mendez, J. Petersson, K. Wallace, A. M. Robinson, R. B. Thakkar, G. D’Haens, Effect of tight control management on Crohn’s disease (CALM): a multicentre, randomised, controlled phase 3 trial. Lancet 390, 2779–2789 (2018).

18. L. Peyrin-Biroulet, W. Reinisch, J. F. Colombel, G. J. Mantzaris, A. Kornbluth, R. Diamond, P. Rutgeerts, L. K. Tang, F. J. Cornillie, W. J. Sandborn, Clinical disease activity, C-reactive protein normalisation and mucosal healing in Crohn’s disease in the SONIC trial. Gut 63, 88–95 (2014).

19. L. Peyrin-Biroulet, W. Sandborn, B. E. Sands, W. Reinisch, W. Bemelman, R. V. Bryant, G. D’Haens, I. Dotan, M. Dubinsky, B. Feagan, G. Fiorino, R. Gearry, S. Krishnareddy, P. L. Lakatos, E. V. Loftus, Jr., P. Marteau, P. Munkholm, T. B. Murdoch, I. Ordas, R. Panaccione, R. H. Riddell, J. Ruel, D. T. Rubin, M. Samaan, C. A. Siegel, M. S. Silverberg, J. Stoker, S. Schreiber, S. Travis, G. Van Assche, S. Danese, J. Panes, G. Bouguen, S. O’Donnell, B. Pariente, S. Winer, S. Hanauer, J. F. Colombel, Selecting Therapeutic Targets in Inflammatory Bowel Disease (STRIDE): Determining Therapeutic Goals for Treat-to-Target. Am. J. Gastroenterol. 110, 1324–1338 (2015).

20. G. Kojouharoff, W. Hans, F. Obermeier, D. N. Mannel, T. Andus, J. Scholmerich, V. Gross, W. Falk, Neutralization of tumour necrosis factor (TNF) but not of IL-1 reduces inflammation in chronic dextran sulphate sodium-induced colitis in mice. Clin. Exp. Immunol. 107, 353–358 (1997).

21. M. Sasaki, S. Bharwani, P. Jordan, T. Joh, K. Manas, A. Warren, H. Harada, P. Carter, J. W. Elrod, M. Wolcott, M. B. Grisham, J. S. Alexander, The 3-hydroxy-3-methylglutaryl-CoA reductase inhibitor pravastatin reduces disease activity and inflammation in dextran-sulfate induced colitis. J. Pharmacol. Exp. Ther. 305, 78–85 (2003).

22. A. Lei, Q. Yang, X. Li, H. Chen, M. Shi, Q. Xiao, Y. Cao, Y. He, J. Zhou, Atorvastatin promotes the expansion of myeloid-derived suppressor cells and attenuates murine colitis. Immunology 149, 432–446 (2016).

23. E. Antoniou, G. A. Margonis, A. Angelou, A. Pikouli, P. Argiri, I. Karavokyros, A. Papalois, E. Pikoulis, The TNBS-induced colitis animal model: An overview. Annals of medicine and surgery (2012) 11, 9–15 (2016).

24. P. S. Dulai, B. S. Boland, S. Singh, K. Chaudrey, J. L. Koliani-Pace, G. Kochhar, M. P. Parikh, E. Shmidt, J. Hartke, P. Chilukuri, J. Meserve, D. Whitehead, R. Hirten, A. C. Winters, L. G. Katta, F. Peerani, N. Narula, K. Sultan, A. Swaminath, M. Bohm, D. Lukin, D. Hudesman, J. T. Chang, J. Rivera-Nieves, V. Jairath, G. Y. Zou, B. G. Feagan, B. Shen, C. A. Siegel, E. V. Loftus, Jr., S. Kane, B. E. Sands, J. F. Colombel, W. J. Sandborn, K. Lasch, C. Cao, Development and Validation of a Scoring System to Predict Outcomes of Vedolizumab Treatment in Patients With Crohn’s Disease. Gastroenterology 155, 687–695 e610 (2018).

25. D. Nummer, E. Suri-Payer, H. Schmitz-Winnenthal, A. Bonertz, L. Galindo, D. Antolovich, M. Koch, M. Buchler, J. Weitz, V. Schirrmacher, P. Beckhove, Role of tumor endothelium in CD4+ CD25+ regulatory T cell infiltration of human pancreatic carcinoma. J. Natl. Cancer Inst. 99, 1188–1199 (2007).

26. H. K. Drescher, A. Schippers, T. Clahsen, H. Sahin, H. Noels, M. Hornef, N. Wagner, C. Trautwein, K. L. Streetz, D. C. Kroy, beta7-Integrin and MAdCAM-1 play opposing roles during the development of non-alcoholic steatohepatitis. J. Hepatol. 66, 1251–1264 (2017).

27. F. Perez-Balderas, S. I. van Kasteren, A. A. Aljabali, K. Wals, S. Serres, A. Jefferson, M. Sarmiento Soto, A. A. Khrapitchev, J. R. Larkin, C. Bristow, S. S. Lee, G. Bort, F. De Simone, S. J. Campbell, R. P. Choudhury, D. C. Anthony, N. R. Sibson, B. G. Davis, Covalent assembly of nanoparticles as a peptidase-degradable platform for molecular MRI. Nature communications 8, 14254 (2017).

28. W. J. Sandborn, S. D. Lee, D. Tarabar, E. Louis, M. Klopocka, J. Klaus, W. Reinisch, X. Hebuterne, D. I. Park, S. Schreiber, S. Nayak, A. Ahmad, A. Banerjee, L. S. Brown, F. Cataldi, K. J. Gorelick, J. B. Cheng, M. Hassan-Zahraee, R. Clare, G. R. D’Haens, Phase II evaluation of anti-MAdCAM antibody PF-00547659 in the treatment of Crohn’s disease: report of the OPERA study. Gut, (2017).

29. E. Kaaru, A. Bianchi, A. Wunder, V. Rasche, D. Stiller, Molecular Imaging in Preclinical Models of IBD with Nuclear Imaging Techniques: State-of-the-Art and Perspectives. Inflamm. Bowel Dis. 22, 2491–2498 (2016).

30. J. L. Tlaxca, J. J. Rychak, P. B. Ernst, P. R. Konkalmatt, T. I. Shevchenko, T. T. Pizarro, J. Rivera-Nieves, A. L. Klibanov, M. B. Lawrence, Ultrasound-based molecular imaging and specific gene delivery to mesenteric vasculature by endothelial adhesion molecule targeted microbubbles in a mouse model of Crohn’s disease. J. Control. Release 165, 216–225 (2013).

31. C. Bachmann, A. L. Klibanov, T. S. Olson, J. R. Sonnenschein, J. Rivera-Nieves, F. Cominelli, K. F. Ley, J. R. Lindner, T. T. Pizarro, Targeting mucosal addressin cellular adhesion molecule (MAdCAM)-1 to noninvasively image experimental Crohn’s disease. Gastroenterology 130, 8–16 (2006).

32. F. Knieling, C. Neufert, A. Hartmann, J. Claussen, A. Urich, C. Egger, M. Vetter, S. Fischer, L. Pfeifer, A. Hagel, C. Kielisch, R. S. Gortz, D. Wildner, M. Engel, J. Rother, W. Uter, J. Siebler, R. Atreya, W. Rascher, D. Strobel, M. F. Neurath, M. J. Waldner, Multispectral Optoacoustic Tomography for Assessment of Crohn’s Disease Activity. N. Engl. J. Med. 376, 1292–1294 (2017).

33. N. Zarghami, A. A. Khrapitchev, F. Perez-Balderas, M. S. Soto, J. R. Larkin, L. Bau, N. R. Sibson, Optimization of molecularly targeted MRI in the brain: empirical comparison of sequences and particles. International journal of nanomedicine 13, 4345–4359 (2018).

34. A. Jefferson, N. Ruparelia, R. P. Choudhury, Exogenous microparticles of iron oxide bind to activated endothelial cells but, unlike monocytes, do not trigger an endothelial response. Theranostics 3, 428–436 (2013).

